# Evidence for the temporal regulation of insect segmentation by a conserved set of developmental transcription factors

**DOI:** 10.1101/145151

**Authors:** Erik Clark, Andrew D. Peel

## Abstract

Long-germ insects, such as the fruit fly *Drosophila melanogaster*, pattern their segments simultaneously, whereas short germ insects, such as the beetle *Tribolium castaneum*, pattern their segments sequentially, from anterior to posterior. While the two modes of segmentation at first appear to be very different, many details of segmentation gene expression are surprisingly similar between long-germ and short-germ species. Collectively, these observations hint that insect segmentation may involve fairly conserved patterning mechanisms, which occur within an evolutionarily malleable spatiotemporal framework. Based on genetic and comparative evidence, we now propose that, in both *Drosophila* and *Tribolium* embryos, the temporal progression of the segmentation process is regulated by a temporal sequence of Caudal, Dichaete, and Odd-paired expression. These three transcription factors are broadly expressed in segmenting tissues, providing spatiotemporal information that intersects with the information provided by periodically-expressed segmentation genes such as the pair-rule factors. However, they are deployed differently in long-germ versus short-germ insects, acting as simple timers in *Drosophila*, but as smooth, retracting wavefronts in *Tribolium*, compatible with either gap gene-based or oscillator-based generation of periodicity, respectively.

## INTRODUCTION

An ancestral, conserved and defining feature of arthropods is their possession of a modular body plan composed of distinct segments serially arrayed along the anterior-posterior (AP) body axis. Over the last few decades, comparative developmental studies have revealed that phylogenetically diverse arthropod species share a conserved network of “segment-polarity” genes such as *engrailed (en)* and *wingless (wg)* (Patel 1994; Damen 2002; Farzana & Brown 2008; Janssen & Budd 2013). These genes, which are expressed in a segmentally-reiterated pattern of stripes, encode transcription factors and signalling molecules that together function to organise and maintain the boundaries between segments, as well as the polarity of anatomical features within them (DiNardo et al. 1994; Perrimon 1994; Sanson 2001). However, these comparative studies have also revealed that, across the phylum, quite disparate developmental strategies are used to generate this conserved segmental pattern (reviewed in Peel et al. 2005; Lynch et al. 2012; Williams & Nagy 2016).

The contrast is especially striking for insect segmentation. Although insects exhibit a highly conserved body plan compared to other arthropod clades (the number and identities of segments being largely invariant), the early stages of insect development vary dramatically between species. The distinction between “long-germ” and “short-germ” embryos is particularly notable (Krause 1939; Sander 1976; Davis & Patel 2002; Liu & Kaufman 2005; Rosenberg et al. 2009).

In long-germ embryos, e.g. those of the fruit fly *Drosophila melanogaster*, almost all segments are patterned during the blastoderm stage (Akam 1987; Nasiadka et al. 2002). (The term “long-germ” derives from the fact that, in this “simultaneous” mode of segment formation, the germ rudiment fills much of the volume of the egg from an early stage of embryogenesis.) We now have a detailed understanding of how the *Drosophila* embryo uses a bespoke set of “stripe-specific” enhancer elements, regulated by maternal and “gap” factors, to rapidly establish a periodic pattern of “pair-rule” gene transcription factor expression (Pankratz & Jäckle 1990; Small & Levine 1991; Schroeder et al. 2011). The pair-rule genes are so-called because their expression, each in a pattern of seven regularly-spaced stripes, reflects a transient double-segment periodicity within the *Drosophila* embryo (Nüsslein-Volhard & Wieschaus 1980; Hafen et al. 1984). The positional information in this pattern is then used, at gastrulation, to establish precise patterns of segment-polarity gene expression (14 stripes each). At the same time as the segment-polarity genes begin to be expressed, several of the pair-rule genes also begin to be expressed segmentally, owing to a rewiring of the pair-rule gene network which results in frequency doubling (Clark & Akam 2016a).

In contrast, short-germ embryos, e.g. those of the beetle *Tribolium castaneum*, have retained the ancestral arthropod condition of patterning their segments in a pronounced sequential sequence from anterior to posterior over the course of embryogenesis (Patel et al. 1994; Patel 1994; Choe et al. 2006; Choe & Brown 2007). (The term “short-germ” derives from the fact that these embryos pattern only their anterior segments at the blastoderm stage, from what is usually a shorter germ rudiment restricted to a posterior-ventral region of the early egg.) The more posterior segments are patterned after gastrulation, in a process that is often coupled to embryo growth.

In *Tribolium*, as well as a species of myriapod, the segmentation process has been shown to involve periodic oscillations of pair-rule gene expression in the posterior of the embryo (Sarrazin et al. 2012; El-Sherif et al. 2012; Brena & Akam 2013). Similar dynamic patterns of pair-rule gene expression have also been found in spiders, crustaceans, and other short-germ insects (for example, Schönauer et al. 2016; Eriksson et al. 2013; Mito et al. 2007). These findings have drawn parallels with vertebrate somitogenesis, which is thought to occur via a “clock and wavefront” mechanism (Cooke & Zeeman 1976; Palmeirim et al. 1997; Oates et al. 2012). Indeed, pair-rule gene orthologs in short-germ arthropods appear to be either components of, or entrained by, a segmentation clock (Stollewerk et al. 2003; Choe et al. 2006; Pueyo et al. 2008). Their striped expression patterns are thus generated by sustained oscillatory dynamics rather than through precise positioning by gap gene orthologs.

The terms “short-germ” and “long-germ” on their own do not adequately describe the true diversity of developmental modes observed within the insect clade, and are perhaps better considered opposite ends of an embryological and/or evolutionary spectrum (Davis & Patel 2002). Individual insect species may fall anywhere on a continuum from extreme “short-germ” to extreme “long-germ” development, depending partly upon the number of segments patterned before versus after gastrulation. Although long-germ development is only found within holometabolous insects, each order within Holometabola contains both shorter-germ and longer-germ species, suggesting that long-germ segmentation has evolved from a short-germ ancestral state several times independently (Davis & Patel 2002; Jaeger 2011). There is also at least one documented case of long-germ segmentation reverting to the short-germ state (Sucena et al. 2014).

These observations argue that evolutionary transitions between the sequential/short-germ and simultaneous/long-germ modes of segment patterning must be relatively straightforward. At first sight, this inference is hard to square with the contrasting patterning strategies involved. However, close inspection of orthologous segmentation gene expression between long-germ and short-germ embryos (Figure 1) reveals striking commonalities, suggesting that these overt differences might mask an underlying mechanistic conservation (Patel et al. 1994; Davis et al. 2001; Damen et al. 2005; Davis et al. 2005; Choe et al. 2006; Janssen, Budd & Damen 2011; Green & Akam 2013). First, as alluded to above, all arthropods use periodically expressed pair-rule gene orthologs to initialise the segment pattern. Second, across all arthropods there is a temporal distinction between the earlier-expressed “primary” pair-rule genes (i.e. *hairy, even-skipped, runt, odd-skipped*, and, in some species, *ftz)* and the later-expressed “secondary” pair-rule genes (i.e. *paired* and *sloppy-paired).* Third, the primary pair-rule genes tend to be initially expressed dynamically, while the secondary pair-rule genes are expressed in stable stripes that prefigure segment-polarity gene expression and static segmental boundaries. Finally, judging by relative expression patterns, the way that the secondary pair-rule genes regulate the segment-polarity genes is likely strongly conserved (Figure 1C, right).

**Fig. 1:**
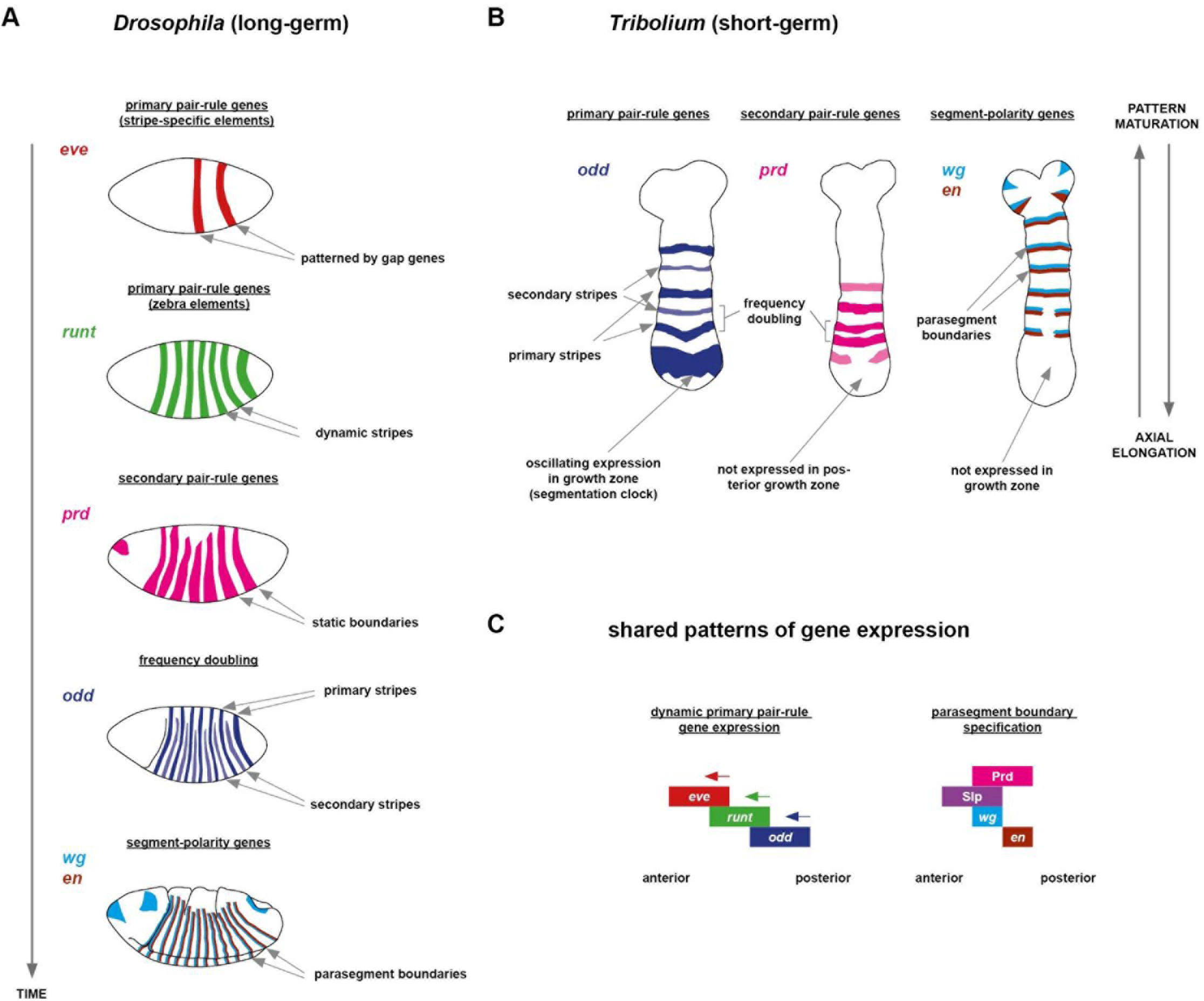
Schematic overview of segment patterning in *Drosophila* versus *Tribolium.* **A:** Overview of pair-rule patterning in *Drosophila.* Top row depicts expression from the *eve* 4+6 reporter element, other rows depict endogenous expression patterns. Time axis runs from early cellularisation (top) to early germband extension (bottom). **B:** Overview of segmentation in *Tribolium.* Embryos are depicted at mid germband extension, but gene expression patterns are essentially similar throughout axial elongation. **C:** Similar relative patterns of gene expression are seen in both long-germ and short-germ species. Left: a dynamic sequence of eve, *runt*, and *odd* expression is seen both *Drosophila* blastoderm and *Tribolium* growth zone (Clark 2017; Choe et al. 2006). Right: the same pattern of secondary pair-rule gene expression is seen upstream of parasegment boundary specification in all arthropods yet studied, including *Drosophila* and *Tribolium* (Green & Akam 2013). Morphological boundaries in the embryo form later, located between the abutting *wg* and *en* domains.

The unexpected similarities between long-germ and short-germ segmentation detailed above are much easier to rationalise in the light of a revised model of *Drosophila* segment patterning, which posits that dynamic shifts of pair-rule stripes are essential for the correct patterning of segment-polarity genes (Clark 2017). With this model, the *Drosophila* pair-rule gene network can easily be modified into a clock and wavefront-type system; depending on initial conditions, the simulated network can broadly recapitulate the patterning dynamics seen in either long-germ or short-germ insects, in both cases giving rise to the same final output pattern of segment-polarity gene expression. The model therefore suggests that long-germ and short-germ modes of segmentation represent alternate tissue-level behaviours of a fundamentally conserved transcriptional network, with similar cellular-level expression dynamics observed in each case. Specifically, the model proposes that the choice between these alternate macroscopic behaviours (simultaneous or sequential segment patterning) is determined by the particular spatiotemporal expression patterns of extrinsic inputs that control the timing of state transitions within the pair-rule network.

The insights from the modelling study rest on two key assumptions/predictions, which can be experimentally examined and tested in short-germ and long-germ insect models. First, there should exist broadly expressed factors that, via their influences on the segmentation network, control the temporal progression of the segmentation process. Two such factors have already been identified: in *Drosophila*, the transcriptional cofactor Odd-paired (Benedyk et al. 1994), which precipitates the transition from double-segment to single-segment periodicity (Clark & Akam 2016a), and, in *Tribolium*, the homeodomain transcription factor Caudal (Schulz et al. 1998; Macdonald & Struhl 1986), which is thought to quantitatively tune the dynamics of the segmentation clock (El-Sherif et al. 2014). Second, evolutionary transitions between segmentation modes should be effected in part by altered expression profiles of these extrinsic “timing factors”, which then drive altered downstream expression of the segmentation genes. Consequently, the spatiotemporal expression of Odd-paired, Caudal, and other timing factors should remain tightly correlated with specific phases of segmentation gene expression across all insect embryos, regardless of whether they exhibit a long-germ or short-germ mode of development.

In this manuscript, we first identify Caudal and Dichaete as additional factors playing important temporal roles in *Drosophila* segmentation. We chose to study these two genes because earlier studies have shown that mutations in these genes affect the segment pattern (Macdonald & Struhl 1986; Russell et al. 1996; Nambu & Nambu 1996), but the precise reasons why have never been clear. We then show that, as predicted, *Tribolium* orthologs of *caudal, Dichaete*, and *odd-paired* are expressed in the same temporal order, and preserve the same correlations with segmentation gene expression as are observed in *Drosophila.* However, while in *Drosophila* these factors are expressed ubiquitously throughout the trunk and thus act as simple timers, in *Tribolium* they provide retracting or advancing wavefronts, and thus could represent the primary source of spatial information within the short-germ segmentation process. Finally, we discuss the significance of these findings for the evolution of segmentation.

## RESULTS

### *caudal, Dichaete*, and *odd-paired* are expressed sequentially during *Drosophila* segmentation

Figure 2 shows the expression of *caudal (cad), Dichaete* (D), and *odd-paired (opa)* transcripts over the course of *Drosophila* segmentation, relative to the expression of the pair-rule gene *odd-skipped (odd)*, which is here used as a marker to show the progressive refinement of segment patterning. We find that all three of these candidate timer genes are widely expressed throughout the trunk at some point during segmentation, consistent with previous descriptions of their expression patterns (Macdonald & Struhl 1986; Russell et al. 1996; Nambu & Nambu 1996; Benedyk et al. 1994). However, the phase of widespread expression spans a different developmental time period in each case, meaning that the segmentation process occurs against a changing background of these transcription factors’ expression.

**Fig. 2:**
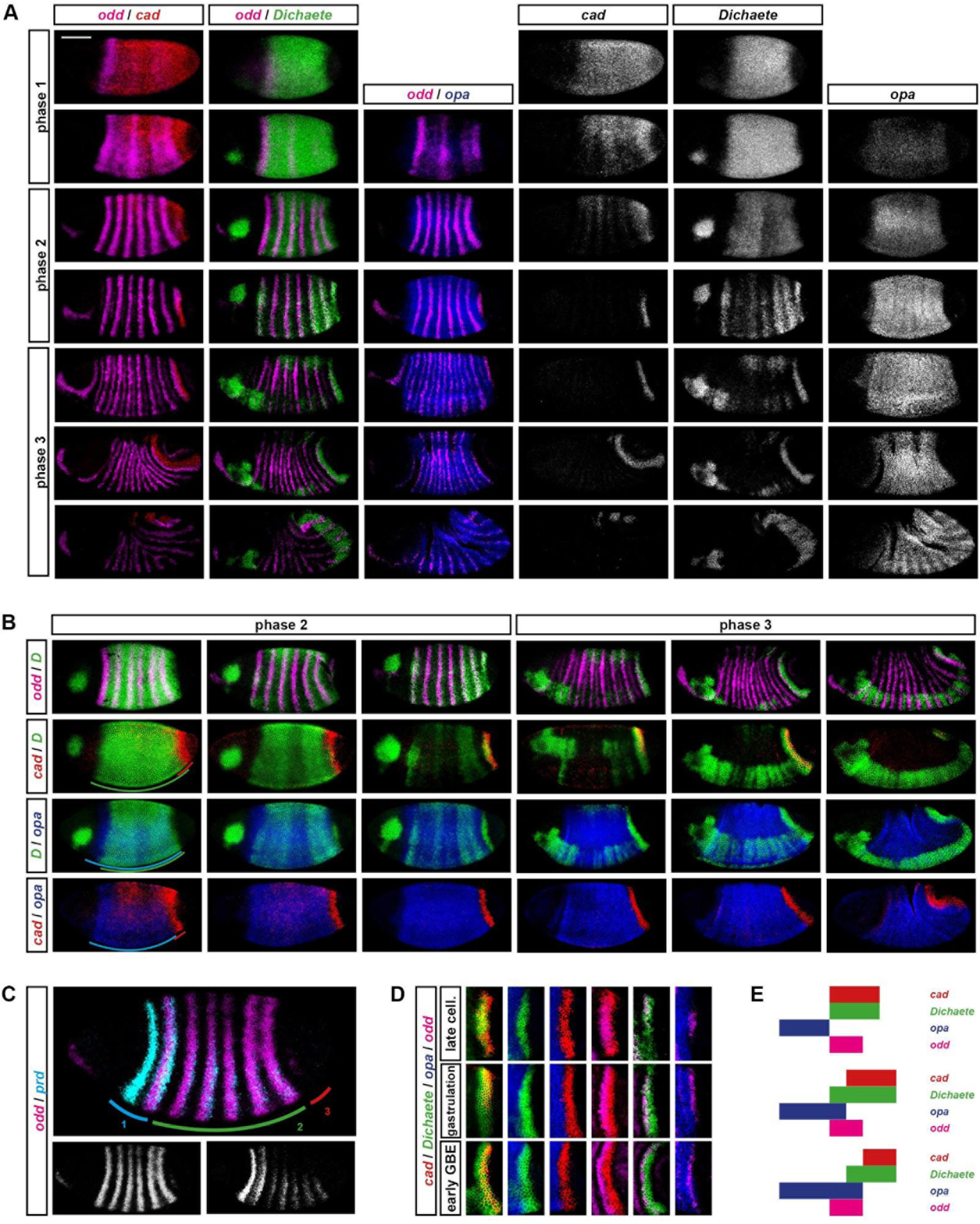
*cad, Dichaete*, and *opa* are expressed dynamically over the course of *Drosophila* segmentation. **A:** Spatiotemporal expression of cad, *Dichaete*, and opa, relative to the pair-rule gene odd. Embryo age increases from top to bottom. “Phase 1” = early cellularisation; “phase 2” = mid-late cellularisation; “phase 3” = gastrulation and early germband extension. Individual channels for cad, *Dichaete*, and *opa* are shown on the right of the false-coloured dual-channel images. All *Drosophila* embryos here and elsewhere are aligned anterior left, dorsal top. Scale bar = 100 μm. The *opa* / *odd* in situs are from Clark and Akam (2016a). **B:** Relative locations of the cad, *Dichaete*, and *opa* expression domains, from mid-cellularisation (early phase 2) onwards. Equivalently-staged *Dichaete/odd* in situs are also shown (top row), for easy cross-referencing with **A.** Embryo age increases from left to right. Annotations in left-most column highlight the relative extent of each domain at mid-cellularisation. **C:** Three distinct regions of pair-rule gene expression timing along the AP axis. The embryo shown is just slightly older than those in the left-most column of **B.** Individual channels for *odd* (left) and *prd* (right) are shown below the false-coloured dual-channel image. **D:** Dynamic gene expression within the tail (region 3). Cropped and rotated enlargements of the *odd* stripe 7 region are shown for three timepoints (equivalent to rows 4-6 in **A** and columns 3-5 in **B).** **E:** Schematic diagram of shifting expression domains within the tail, based on the data in **D.** The anterior boundary of *odd* stripe 7 is assumed to be static (see Figure S1 legend).

Following a staging scheme introduced in a recent study of pair-rule gene regulation (Clark & Akam 2016a), pair-rule gene expression can be divided into three broad phases, spanning from early cellularisation to early germband extension (Figure 2A). During phase 1 (early cellularisation), individual stripes of primary pair-rule gene expression are established in an ad hoc manner by stripe-specific enhancer elements, resulting in irregular/incomplete periodic patterns. During phase 2 (mid- to late-cellularisation), pair-rule factors cross-regulate each other through “zebra” elements, resulting in regular stripes of double-segment periodicity, and the secondary pair-rule genes turn on in the trunk. During phase 3 (gastrulation onwards), the regulatory network changes and the expression patterns of some of the pair-rule genes (including *odd*) transition to single-segment periodicity.

*cad* and *Dichaete* are both expressed prior to cellularisation and continue to be expressed during phase 1, meaning that the periodic patterns of the primary pair-rule genes emerge in the context of Cad and Dichaete activity. (Indeed, both Cad and Dichaete have been found to regulate stripe-specific elements: La Rosée et al. 1997; Häder et al. 1998; Ma et al. 1998; Fujioka et al. 1999). *cad* expression retracts to a narrow posterior domain early in phase 2, while *Dichaete* expression persists but gradually fades, passing through a transient phase of periodic modulation. Since the dynamics of Cad and Dichaete protein expression appear to closely reflect their respective transcript patterns (Macdonald & Struhl 1986; Ma et al. 1998), most of phase 2 is likely to be characterised by Dichaete activity alone. *(opa* expression builds up during this time, but owing to the length of the *opa* transcription unit, Opa protein levels are likely to be absent or low (Benedyk et al. 1994).) Finally, phase 3 is characterised by strong *opa* expression, while *Dichaete* becomes expressed in a narrow posterior domain and throughout the neuroectoderm. The widespread changes in pair-rule gene expression that occur during this time are now known to be triggered by Opa activity (Clark & Akam 2016a).

### *caudal, Dichaete*, and *odd-paired* expression spatially correlates with segmentation timing along the anteroposterior axis

In addition to this general temporal progression within the trunk, there are also interesting spatial correlations with the segmentation process (Figure 2B-D). Although *Drosophila* segmentation is generally described as “simultaneous”, the AP axis can actually be divided into three distinct regions, which undergo segment patterning at slightly different times (Figure 2C; Surkova et al. 2008). In region 1 (near the head-trunk boundary, encompassing *odd* stripe 1 and *paired (prd)* stripes 1 and 2), both primary and secondary pair-rule genes are expressed very early, and head-specific factors play a large role in directing gene expression (Schroeder et al. 2011; Andrioli et al. 2004; Chen et al. 2012). In region 2 (the main trunk, encompassing *odd* stripes 2-6 and *prd* stripes 3-7), primary pair-rule genes turn on at phase 1, secondary pair-rules turn on during phase 2, and segment-polarity genes turn on at gastrulation. Finally, in region 3 (the tail, encompassing *odd* stripe 7 and *prd* stripe 8), the expression of certain pair-rule genes is delayed, while segment-polarity gene expression emerges only during germband extension (Kuhn et al. 2000). This region of the fate map eventually gives rise to the “cryptic” terminal segments A9-A11 (Campos-Ortega & Hartenstein 1985; Kuhn et al. 1995).

Suggestively, the boundaries of these three regions correspond to expression boundaries of our candidate timing factors at mid-cellularisation (Figure 2B, left). Region 1 never expresses *cad* or *Dichaete*, but does express *opa*, whose expression domain extends further anterior than those of the other two genes. Region 2 corresponds to the early broad domain of *Dichaete* expression, which extends from just behind *odd* stripe 1 to just behind *odd* stripe 6 (the *opa* expression domain shares the same posterior boundary). Finally, region 3 is characterised by *cad* expression, which extends further posterior than the other two genes, and persists in this posterior region. The expression profiles of cad, *Dichaete*, and *opa* thus correlate with the current state of the segmentation process across space, as well as over time.

Region 3 is particularly intriguing, because as *odd* stripe 7 begins to be expressed, the cells also start to express *Dichaete* (Figure 2B), and as gastrulation proceeds, the cells begin to express *opa* (Figure 2D), thus recapitulating the same temporal sequence that occurred earlier in region 2. Interestingly, we noticed that the expression boundaries of cad, *Dichaete*, and *opa* all shift posteriorly relative to *odd* stripe 7 over time (Figure 2E; for evidence that these are “true” kinematic shifts, see Figure S1).

Given 1) that Opa activity triggers many of the pair-rule gene expression changes observed at gastrulation, 2) that proper segment patterning is also dependent on Cad and Dichaete activity, and 3) that the temporal progression of *cad/Dichaete/opa* expression is correlated with the temporal progression of the segmentation process, it is possible that all three genes provide important temporal regulatory information to the pair-rule system. We therefore investigated whether Cad and Dichaete, like Opa, show regulatory effects on the *Drosophila* pair-rule gene network.

### Caudal controls the onset of *paired* expression

The clearance of *cad* transcript during cellularisation occurs in a distinctive spatiotemporal pattern: expression is lost from anterior regions before posterior regions, and from ventral regions before dorsal regions. A similar spatiotemporal pattern is seen at the protein level (Macdonald & Struhl 1986), suggesting that Cad protein turns over rapidly, and that *cad* transcript patterns therefore provide a reasonable proxy for Cad protein dynamics during this period.

Figure 3 shows that this retraction of *cad* expression is tightly spatiotemporally associated with the emergence of the *prd* stripes. *prd* stripe 1 is established during early cellularisation under the control of a stripe-specific element (Schroeder et al. 2011) and is located just anterior to the broad *cad* domain in the trunk. *prd* stripe 2 follows and then stripes 3-7 emerge during mid-cellularisation, appearing with a marked anterior-to-posterior and ventral-to-dorsal progression that is the inverse of the retracting *cad* domain. Finally, *prd* stripe 8 (asterisk in Figure 3) emerges towards the end of cellularisation, as the *cad* domain in the tail retracts posteriorly (see Figure S1).

**Fig. 3:**
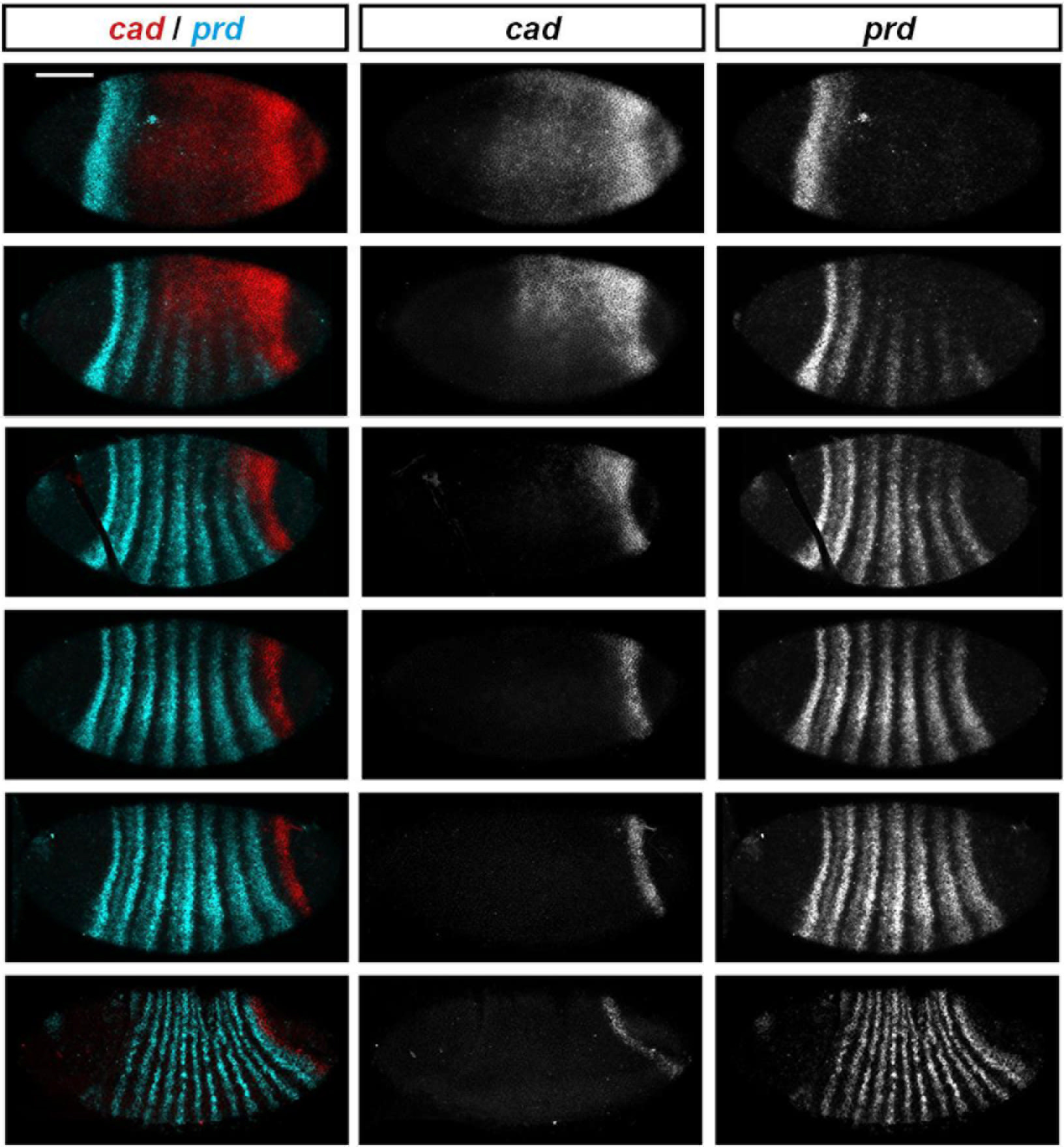
*cad* and *prd* exhibit anti-correlated spatiotemporal dynamics during cellularisation. Embryo age increases from top (early cellularisation) to bottom (gastrulation). Arrowheads mark the posterior border of *prd* stripe 7; asterisk marks *prd* stripe 8. Scale bar = 100 μm.

Setting aside the periodic nature of the *prd* output, *cad* and *prd* thus exhibit strikingly complementary patterns of expression throughout the entire segmentation process, suggesting that Cad might repress *prd* and thus control the time of onset of its expression. To test this, we examined the emergence of the *prd* stripes in *cad* zygotic mutant embryos (Figure 4). Note that we chose to analyse *cad* zygotic mutants, which retain maternal *cad* expression and undergo largely normal segment patterning, rather than complete *cad* nulls, because Cad has pleiotropic effects on the segmentation process, and we wanted to be able to distinguish a possible direct effect on *prd* expression from expected indirect effects stemming from perturbations to early gap gene expression (reviewed in Jaeger 2011).

**Fig. 4:**
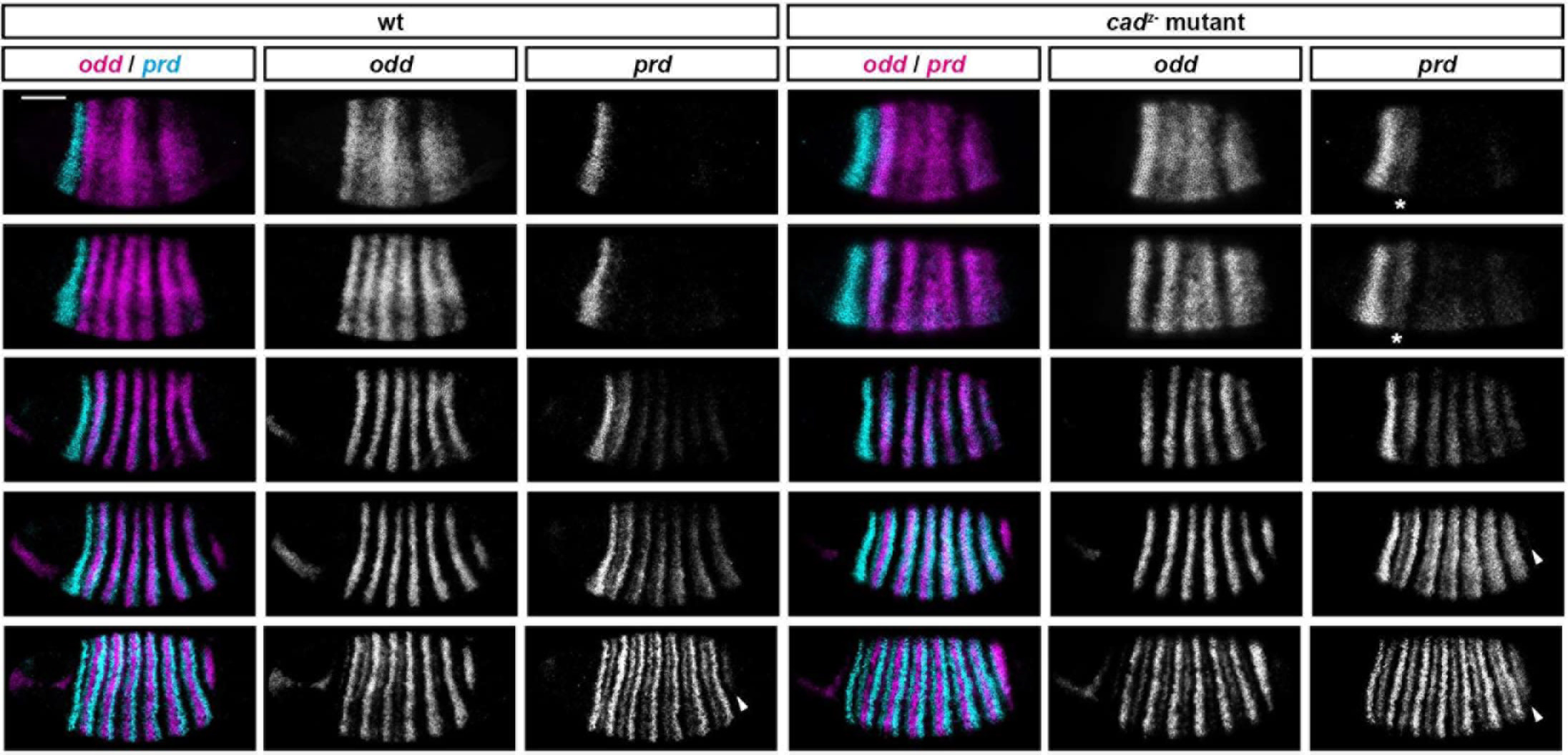
The spatiotemporal onset of *prd* expression is regulated by Cad. *prd* expression is shown in wild-type embryos and *cad* zygotic mutants, using *odd* as a staging marker. (Note the characteristic development of *odd* expression within the head, which is necessarily independent of Cad levels.) Mutant embryos are conservatively staged: note that embryos in rows 2 and 3 are slightly younger than their wild-type counterparts. Asterisks indicate precocious *prd* stripe 2 expression; arrowheads point to emerging *prd* stripe 8 expression. Note the ectopic *prd* expression throughout the trunk in the second mutant embryo down, and the even intensity of the *prd* stripes along the DV axis in the third and fourth mutant embryos down. Scale bar = 100 μm.

Consistent with the proposal that Cad represses *prd* expression, we found that *prd* expression is precocious and more intense in *cad* zygotic mutants. In addition, the stripes turn on simultaneously along the dorsoventral axis rather than in a ventral to dorsal progression, correlating with the absence of the spatially-patterned zygotic Cad domain. (Note, however, that the *prd* stripes remain less intense in dorsal regions of the embryo than elsewhere, indicating that some of the remaining regulators of *prd* expression must be dorsoventrally modulated.)

In contrast to the marked effects on early *prd* expression, other aspects of segmentation gene expression are largely unperturbed in *cad* zygotic mutant embryos, aside from a modest posterior expansion of the fate map (asterisks in Figure S2B). For example, there are only very subtle spatiotemporal changes to the *odd, slp*, and *en* stripes (Figure 4 and Figure S2), and late *prd* expression is also fairly normal, with stripe splitting (an effect of Opa activity) occurring at the same time as in wild-type (Figure 4). These observations indicate that the temporal regulatory information provided by Cad is at least partially decoupled from the general progression of the segmentation process in *Drosophila.*

### Dichaete influences the topology of the early pair-rule network

The cuticles of *Dichaete* null larvae exhibit segmentation defects of varying severity, often involving several segment fusions and/or deletions (Russell et al. 1996; Nambu & Nambu 1996). These defects are not randomly distributed, but tend to affect particular segments (e.g. loss of the A2 and A8 denticle belts, narrowing of the A3 belt, and fusions between A4-A5 and A6-A7). Since gap gene expression is normal in *Dichaete* mutants but pair-rule gene expression is perturbed (Russell et al. 1996), Dichaete is thought to directly regulate pair-rule gene expression, consistent with high levels of Dichaete binding around pair-rule gene loci (Macarthur et al. 2009; Aleksic et al. 2013). However, the specific aetiology of the *Dichaete* mutant phenotype and the reason for its variable severity are not known.

At least some of Dichaete’s regulatory effects are mediated by stripe-specific elements (Ma et al. 1998; Fujioka et al. 1999). However, Dichaete might also act on pair-rule gene “zebra” elements (which take inputs from other pair-rule genes), and thereby influence pair-rule gene cross-regulation. To investigate this possibility, we examined whether any of the interactions within the early (phase 2) pair-rule gene network (Clark & Akam 2016a; Clark 2017) seemed to be perturbed in *Dichaete* mutants.

Five of the pair-rule genes are patterned by other pair-rule genes during phase 2: *runt, ftz, odd, prd*, and slp. Based on double in situ data, the regulatory logic governing ftz, odd, and *slp* expression appears to be normal in *Dichaete* mutant embryos: despite the irregularity of the expression patterns, sharp expression boundaries between repressors and their targets are still observed at mid-cellularisation (Figure S3A). (In other words, even though both the regulatory inputs and transcriptional outputs of each gene are clearly perturbed, normal input-output relationships appear to be preserved.) Note that, in contrast to previous descriptions of *Dichaete* mutants (Russell et al. 1996), we find that pair-rule gene expression patterns are fairly consistent across different embryos, as long as the embryos are similarly-staged.

Surprisingly, we noticed that *odd* is expressed almost ubiquitously throughout the trunk during early cellularisation (phase 1), even though it resolves into a fairly normal pair-rule pattern as cellularisation proceeds (Figure S3B). This ectopic expression looks very similar to the expression pattern of *odd* in *hairy* mutant embryos (Figure S3B), indicating that the effect may be mediated by a functional interaction between Dichaete and Hairy. However, any such interaction would have to be both temporary and target-specific: temporary because Hairy is evidently still able to repress *odd* normally during phase 2, and target-specific because no similar ectopic expression is seen for ftz, which also requires repression from Hairy during phase 1 (Figure S3C).

In contrast to ftz, odd, and slp, the transcriptional outputs of *runt* and *prd* in *Dichaete* mutants are perturbed in ways that indicate significant deviations from their wild-type regulatory logic (Figure 5; Figure 6). The expression patterns of these two genes markedly overlap with those of genes that would normally repress them, demonstrating that these particular regulatory interactions are crucially dependent upon Dichaete activity.

**Fig. 5:**
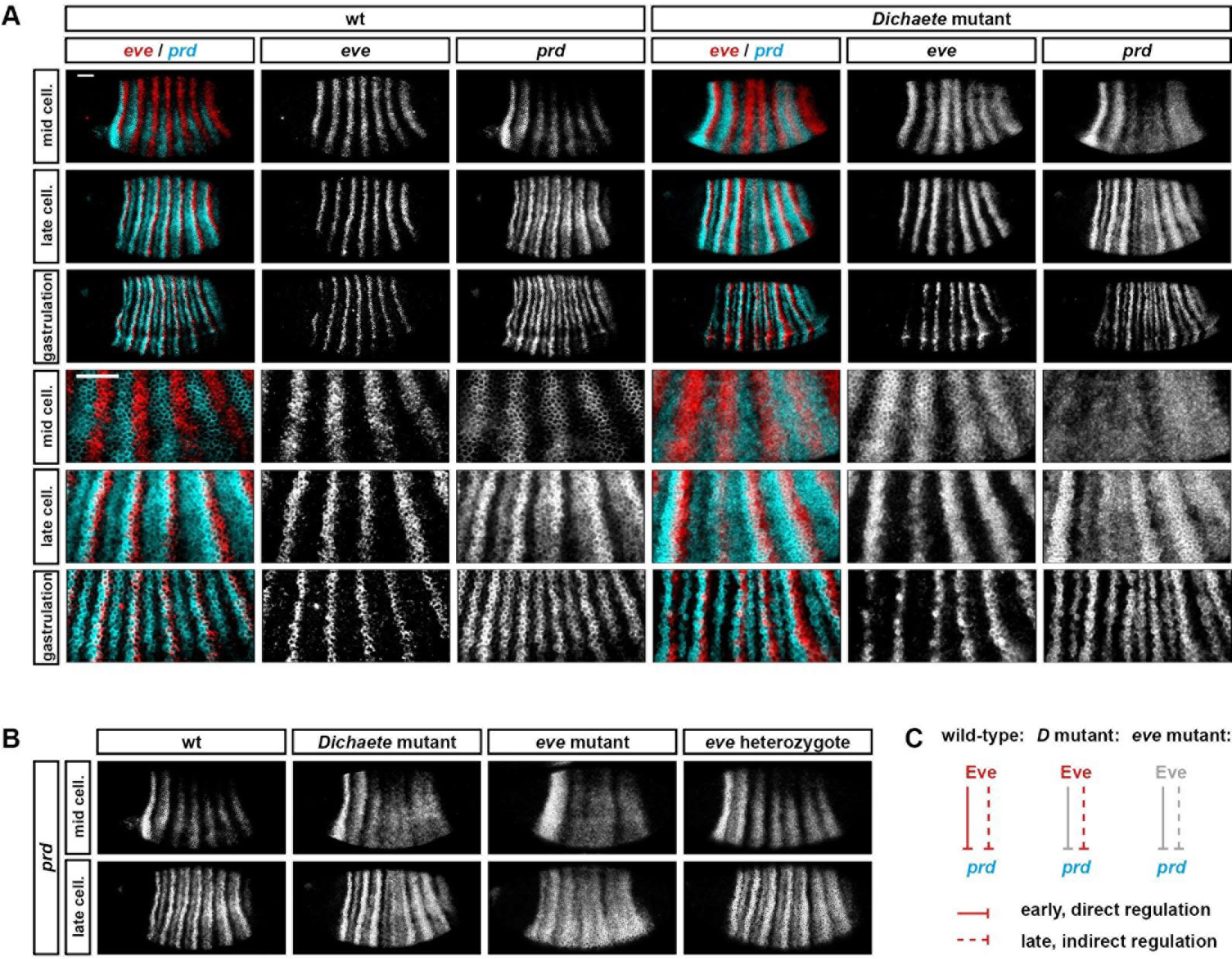
Eve requires Dichaete in order to repress prd. **A:** Expression of *prd* relative to *eve* in wild-type and *Dichaete* mutant embryos, at mid-cellularisation, late cellularisation, and gastrulation. Single channels are shown to the right of each false-coloured dual-channel image, and enlarged views of the trunk region are shown below the whole embryo views. Periodicity of early *prd* expression is lost in the *Dichaete* mutants, but the timing and DV patterning of expression remain normal. Note that *prd* and *eve* stripes overlap at later stages in both wild-type and mutant embryos. Scale bars = 50 μm. **B:** *prd* expression at mid-cellularisation and late-cellularisation, in a variety of genetic backgrounds. A similar loss of *prd* periodicity within the trunk is seen in *Dichaete* and *eve* mutant embryos at mid-cellularisation, but patterning subsequently diverges. **C:** Interpretation of *prd* regulation in **B.**

**Fig. 6:**
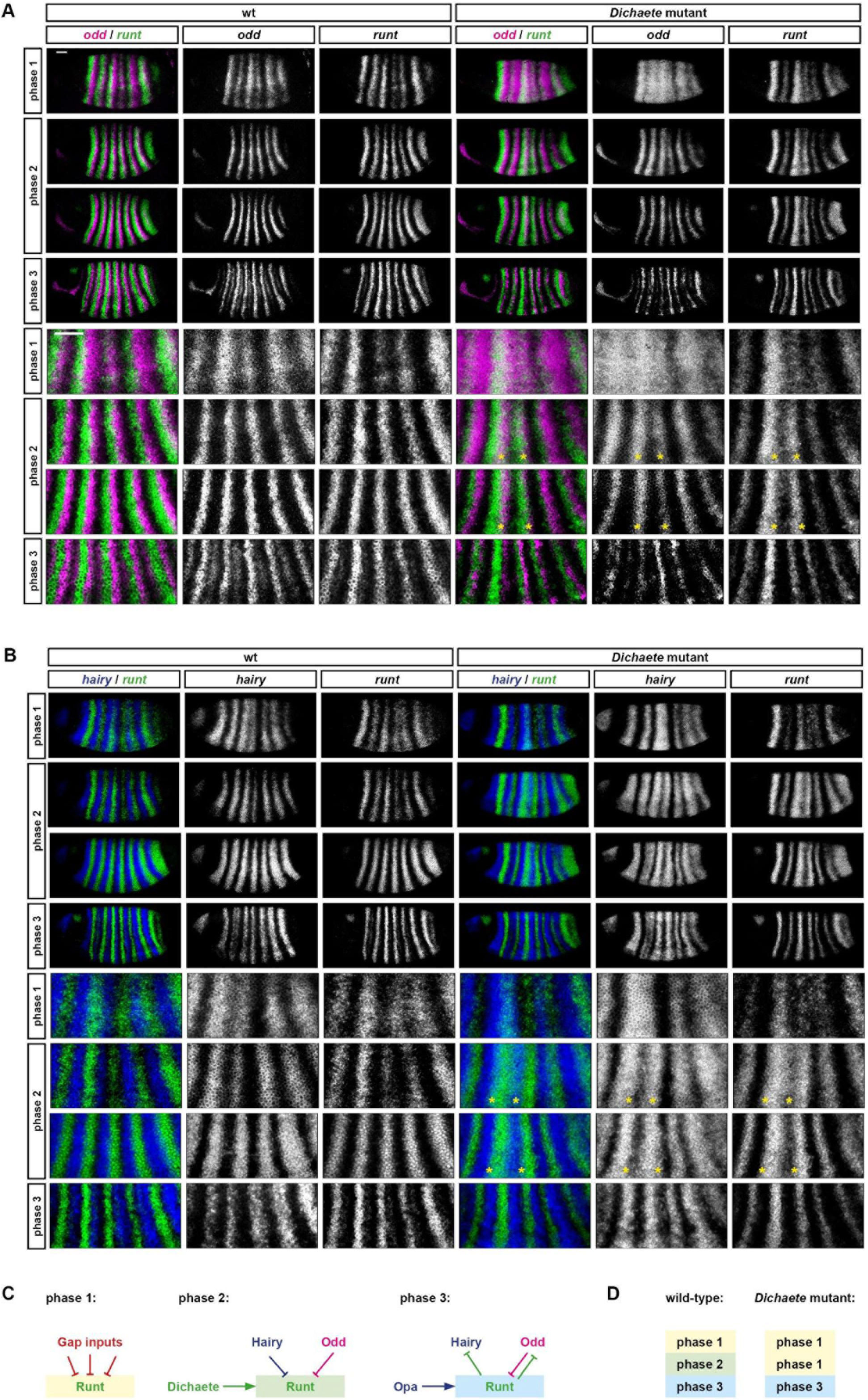
*runt* is not patterned by pair-rule inputs in *Dichaete* mutant embryos. **A, B:** Expression of *runt* relative to *odd* **(A)** and *hairy* **(B)** in wild-type and *Dichaete* mutant embryos, over the course of cellularisation. Single channels are shown to the right of each false-coloured dual-channel image, and enlarged views of the trunk region are shown below the whole embryo views. *runt* expression continues to overlap with *odd* and *hairy* expression throughout phase 2 in *Dichaete* mutants, but not wild-type. Asterisks indicate extensive regions of coexpression associated with *runt* stripes 3 and 4. (Note that narrow strips of overlapping expression in wild-type embryos are caused by the dynamic nature of the pair-rule stripes, which shift anteriorly across the tissue until the beginning of phase 3, causing protein domains to lag slightly behind transcript domains.) Scale bars = 50 μm. **C:** Schematic model of *runt* regulation over the course of segmentation. Only the cross-regulatory interactions between *hairy, odd*, and *runt* are included. **D:** Interpretation of *runt* regulation in **A** and **B.**

*prd* shows the most dramatic expression changes (Figure 5). The early *prd* stripes are normally patterned by direct repression from Eve – they broaden somewhat in *eve* heterozygotes, and fuse into a largely aperiodic expression domain in *eve* mutant embryos (Figure 5B; Baumgartner & Noll 1990; Clark 2017). Although the *eve* stripes are still present in *Dichaete* mutant embryos (Figure 5A), early *prd* expression shows almost as great a loss of periodicity as observed in *eve* mutant embryos, suggesting that the repression of *prd* by Eve requires Dichaete activity. This apparent cofactor role for Dichaete must be target-specific, since other Eve targets are repressed normally in *Dichaete* mutant embryos (see above). Note that the repression of *prd* by Eve is restricted to earlier phases of segmentation in wild-type embryos, and that its temporary nature is likely to be important for segment patterning (Fujioka et al. 1995; Clark 2017). Our results now suggest a mechanistic explanation for the temporal modulation of this regulatory interaction: the loss of Dichaete expression over the course of cellularisation.

Unlike prd, *runt* is still expressed in seven stripes in *Dichaete* mutants, but these stripes are very irregular (Figure 6). Stripes 3 and 7 are broadened compared to wild-type, while stripes 2, 5, and 6 are narrow and fairly weak. In wild-type embryos, the regular stripes of *runt* seen at mid-cellularisation (phase 2) are defined by repression from Hairy at their anteriors, and repression from Odd at their posteriors (Figure 6C; Klingler & Gergen 1993; Saulier-Le Dréan et al. 1998; Clark & Akam 2016a), resulting in sharply abutting expression domains (Figure 6A,B left). However, *runt* expression in *Dichaete* mutant embryos tends to overlap with the stripes of both *hairy* and *odd* (Figure 6A,B right), indicating that this cross-repression is either ineffective or absent. Given the weakened, although still periodic, expression of *runt* in *Dichaete* mutants, combined with the apparent failure to respond to pair-rule inputs, we think that the phase 2 expression we see may derive mainly from *runt’s* stripe-specific elements, rather than mainly from *runt*’s zebra element as in wild-type (Figure 6D). This inference implicates Dichaete as an activator of the zebra element (Figure 6C).

Later aspects of both *runt* and *prd* expression then appear fairly normal in *Dichaete* mutant embryos – as might be expected since Dichaete activity would be largely absent from the trunk during these stages even in wild-type embryos. The *runt* stripes sharpen, and become complementary with those of both *odd* and *hairy*, events that are presumably triggered by known, Opa-dependent regulatory interactions taking effect (Figure 6C; Clark & Akam 2016a). *prd* expression first becomes spatially modulated, and then transitions to a segmental pattern as normal. The initial expression changes are probably regulated by Runt (compare Figure 6), accounting for their irregularity, while stripe splitting is another Opa-dependent effect (Baumgartner & Noll 1990).

However, although pair-rule gene regulatory logic thus appears completely wild-type during these later stages, the resulting expression patterns are of course abnormal, owing to the previous misregulation of key genes. Figure S4 shows how these early patterning irregularities are carried forward, eventually resulting in segment boundary defects that prefigure the cuticle defects observed at the end of development. (Specifically, irregular *runt* stripes lead to abnormal widths and spacing of the *slp* stripes, something which then has knock-on effects for the final pattern of segment-polarity fates.) We have therefore shown that the *Dichaete* segmentation phenotype follows in a predictable manner from abnormal patterns of pair-rule gene expression laid down at the blastoderm stage, illustrating the importance of precise early stripe phasing for producing a viable embryo.

### *Tc-caudal, Tc-Dichaete*, and *Tc-odd-paired* provide staggered wavefronts that likely regulate *Tribolium* segmentation

We have shown above that Cad, Dichaete, and Odd-paired all have important, and temporally distinct, influences on the *Drosophila* pair-rule gene network. Given that pair-rule gene orthologs carry out segment patterning in both long-germ and short-germ insects, we wondered whether the roles of these “timing factors” might also be broadly conserved. We therefore examined the expression of their orthologs, *Tc-cad, Tc-Dichaete*, and *Tc-opa*, in the short-germ beetle *Tribolium castaneum.* The expression patterns of *Tc-cad* and *Tc-Dichaete* have been described previously (Schulz et al. 1998; Oberhofer et al. 2014), but only for a small number of stages, affording only a limited view into their spatiotemporal dynamics over the course of segmentation. *opa* expression has been previously looked at in spiders and myriapods (Damen et al. 2005; Janssen, Budd, Prpic, et al. 2011), but to our knowledge has not been examined in non-dipteran insects.

Figure 7 shows staged expression of *Tc-cad, Tc-Dichaete*, and *Tc-opa*, all relative to a common marker, *Tc-wg*, over the course of germband extension. A more extensive set of stages is shown in Figure S5, and direct comparisons between *Tc-cad/Tc-Dichaete, Tc-cad/Tc-opa*, and *Tc-Dichaete/Tc-opa* are shown in Figure S6.

**Fig. 7.**
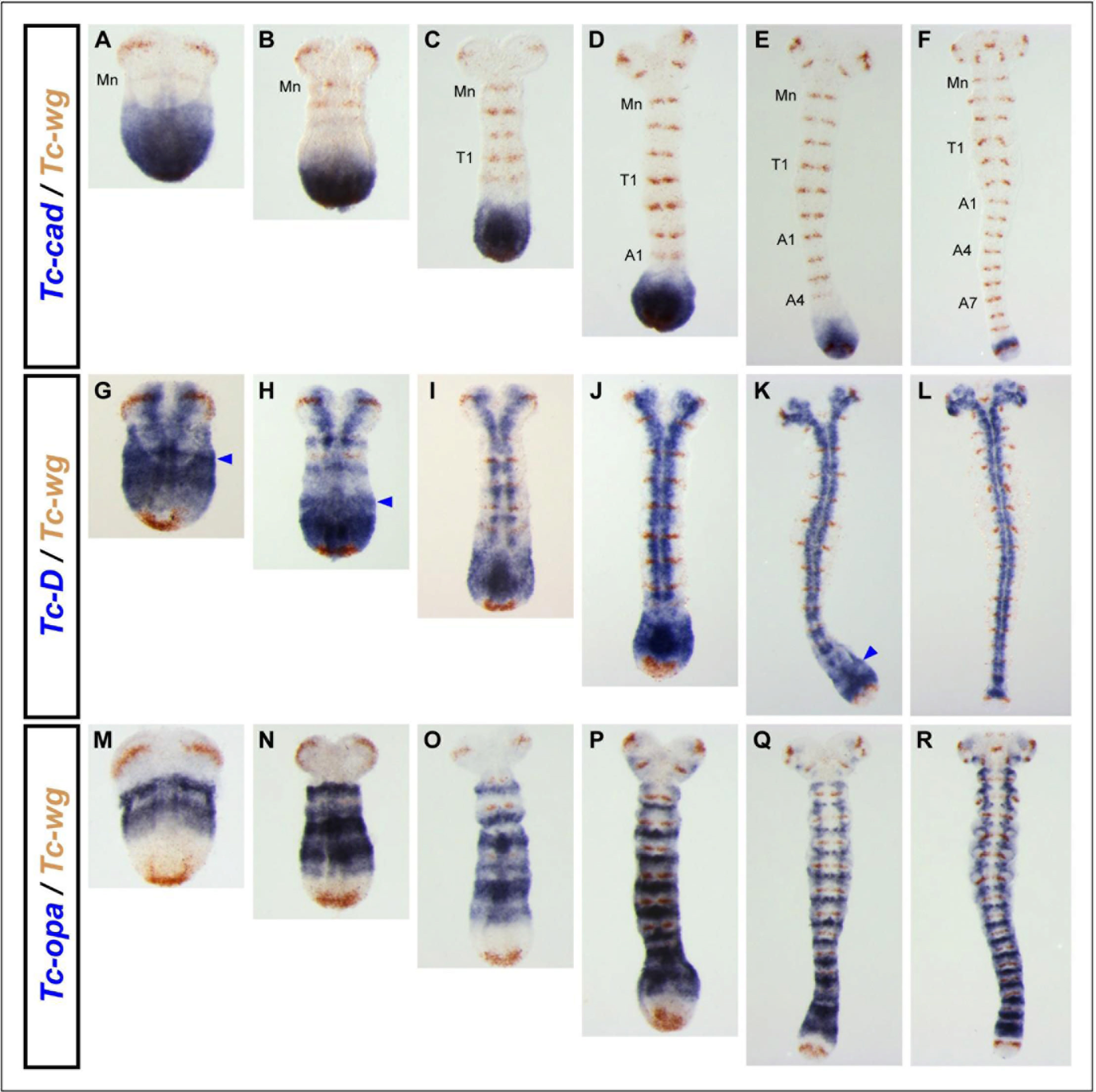
Expression of *Tc-cad, Tc-Dichaete* & *Tc-opa* relative to a common segment marker, *Tc-wg*, in *Tribolium castaneum* germband stage embryos. **(A-F).** Double in situ hybridization for *Tc-cad* (blue) and *Tc-wg* (brown) in embryos of increasing age from left **(A)** to right **(F).** The mandibular (Mn), prothoracic (T1), 1^st^ abdominal (A1), 4^th^ abdominal (A4) & 7th abdominal (A7) stripes of *Tc-wg* expression have been labeled. **(G-L).** As for panels A-F, but this time double in situ hybridization for *Tc-Dichaete* (blue) and *Tc-wg* (brown). Blue arrowheads point to a stripe of *Tc-Dichaete* expression that has formed, or is forming, anterior to the strong posterior expression domain (see text for further details). **(M-R).** As for **A-F,** but this time double in situ hybridization for *Tc-opa* (blue) and *Tc-wg* (brown). Panels in the same column have been stage matched using *Tc-wg* expression patterns. Note how the *Tc-cad, Tc-Dichaete*, and *Tc-opa* expression patterns shift posteriorly relative to *Tc-wg* expression as embryos elongate, and the remarkable consistency of these posterior expression domains during the process of germband elongation. Consult Fig. S5 for a more extensive range of stages, which are presented in a manner that aids comparison between *Tc-cad, Tc-Dichaete* & *Tc-opa* expression patterns, rather than variation across developmental stages.

*Tc-cad* is continuously expressed in the growth zone, resulting in a persistent posterior domain that gradually shrinks over time. *Tc-cad* therefore turns off in presegmental tissues just as they emerge from the anterior of the growth zone, and well before the *Tc-wg* stripes turn on.

*Tc-Dichaete* is also broadly expressed within the growth zone, but is excluded from the most posterior tissue, turning on slightly anterior to the terminal *Tc-wg* domain. The growth zone expression of *Tc-Dichaete* extends slightly further to the anterior than that of *Tc-cad*, turning off just before the *Tc-wg* stripes turn on. *(Tc-Dichaete* expression anterior to the *Tc-cad* domain tends to be at lower levels and/or periodically modulated.) In a subset of embryos we observe a separate stripe of *Tc-Dichaete* expression, anterior to growth zone domain, which clearly encompasses lateral ectodermal cells (Figure 8; Figure S6, A-L). *Tc-Dichaete* expression later transitions into persistent expression within the neuroectoderm, with expression now absent from the more lateral ectodermal regions.

**Fig. 8.**
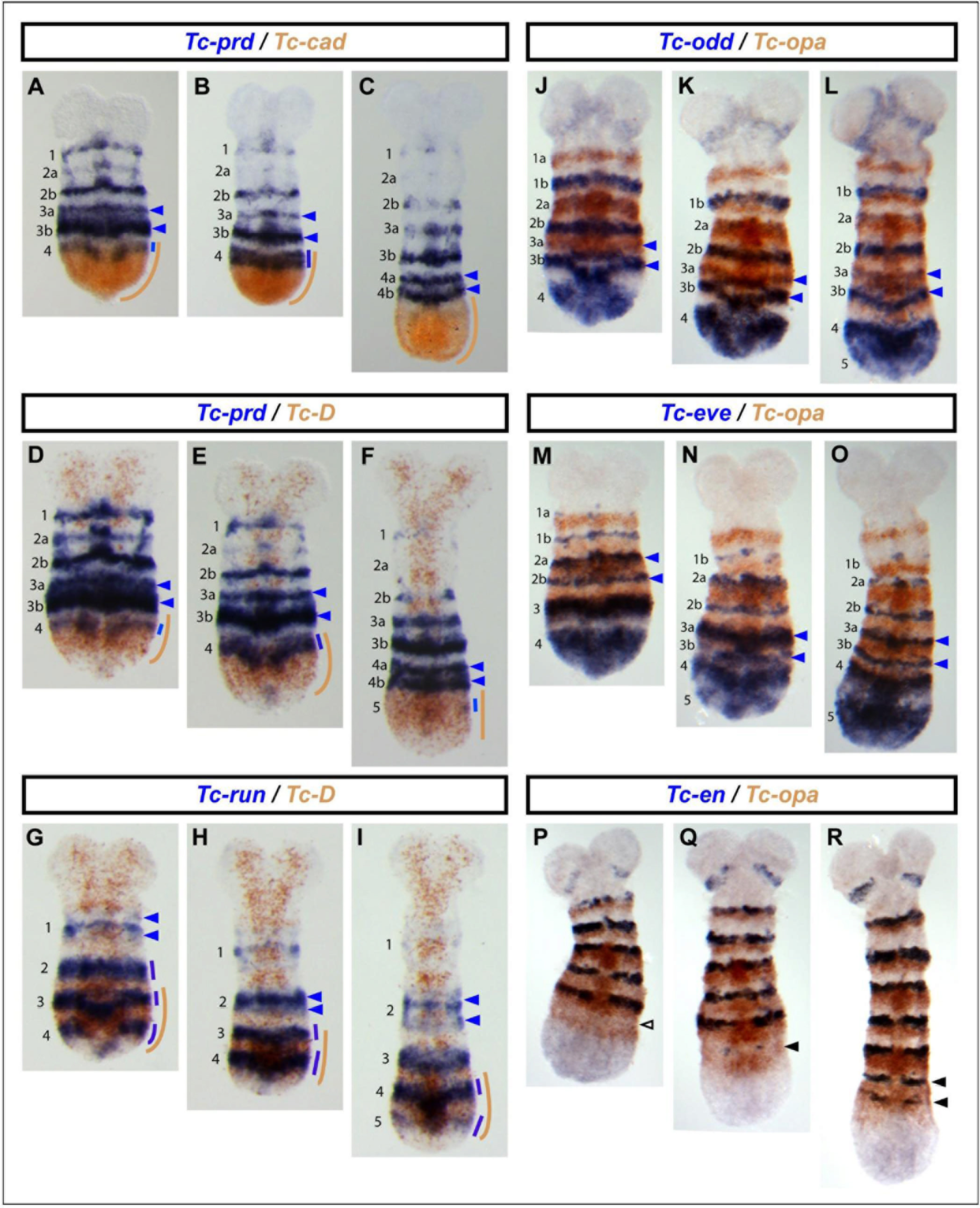
Expression of *Tc-cad, Tc-Dichaete* & *Tc-opa* relative to selected segmentation genes in *Tribolium castaneum* germband stage embryos. **(A-C).** Double in situ hybridization for *Tc-prd* (blue) and *Tc-cad* (brown) on three embryos that span the formation and splitting of the 4^th^ *Tc-prd* primary pair-rule stripe. **(D-F).** As for panels **A-C,** but this time double in situ hybridization for *Tc-prd* (blue) and *Tc-Dichaete* (brown). Note how the primary pair-rule stripe of *Tc-prd* expression forms at the anterior of the *Tc-cad* domain **(A, B)** and the anterior of the posterior-most *Tc-Dichaete* domain **(D, E),** but only splits to form segmental stripes (labeled 4a & 4b) anterior to these domains **(C, F). (G-I).** Double in situ hybridization for *Tc-run* (blue) and *Tc-Dichaete* (brown) on three embryos that span the formation of the 4^th^ and 5^th^ *Tc-run* primary pair-rule stripes, plus splitting of the 2^nd^ *Tc-run* primary pair-rule stripe. Note how *Tc-run* stripe splitting occurs anterior to the posterior-most *Tc-Dichaete* domain. **(J-L).** Double in situ hybridization for *Tc-odd* (blue) and *Tc-opa* (brown) on three embryos that span the formation of the 4^th^ and 5^th^ *Tc-odd* primary pair-rule stripes and the resolution of the 3^rd^ *Tc-odd* primary pair-rule stripe into two segmental stripes (labeled 3a & 3b). **(M-O).** As for **J-L,** but this time double *in situ* hybridization for *Tc-eve* (blue) and *Tc-opa* (brown). Note how segmented stripes of *Tc-eve* and *Tc-odd* expression resolve within the *Tc-opa* domain **(K, L, N, O).** In **A-O,** blue arrowheads mark *Tc-prd, Tc-run, Tc-odd* & *Tc-eve* segmental stripes that have recently resolved, or are in the process of resolving. **(P-R).** Double in situ hybridization for *Tc-en* (blue) and *Tc-opa* (brown) on three embryos that span the formation of the T3 and A1 *Tc-en* segmental stripes. Note how the forming *Tc-en* stripes (solid black arrowheads in **Q, R)** appear within the *Tc-opa* domain, but in a stripe-shaped region where *Tc-opa* expression is already clearing (empty black arrowhead in **P).** Colour-coded lines on the right-hand side of the embryos indicate our interpretations of the expression patterns in **A-I.** For a more detailed description of these expression patterns, consult the main text and supplementary figures 7-10.

Finally, *Tc-opa* is absent from the posterior half of the growth zone, but is expressed in a broad posterior domain starting in the anterior growth zone and extending anteriorly far enough to overlap nascent *Tc-wg* stripes. The intensity of expression in this domain is periodically-modulated, with *Tc-opa* expression transitioning more anteriorly into relatively persistent segmental stripes. The segmental stripes cover the central third of each segment, lying posterior to each *Tc-en* stripe (Figure 8R).

The most striking aspect of these expression patterns is that while they are obviously dynamic, retracting posteriorly as the embryo elongates, they appear remarkably consistent from one stage to the next: the posterior expression domains of each of the three genes retain the same relationship to the gross morphology of the embryo, and to each other, at each timepoint. This means that each cell that starts off within the growth zone will at some point experience a temporal progression through the three transcription factors, similar to that experienced by cells within the *Drosophila* trunk over the course of cellularisation and gastrulation (compare Figure 2). For most of the cells that contribute to the *Tribolium* trunk, this sequence is likely to start with Cad+Dichaete, transit through Dichaete+Opa, and end with Opa alone (compare Figure 7 against the *Tribolium* lineage tracing experiments from Nakamoto et al. 2015).

We also compared the expression domains of *Tc-cad, Tc-Dichaete*, and *Tc-opa* to the expression patterns of a number of *Tribolium* segmentation genes (Figure 8; and more extensive developmental series in Figures S7–S10). Strikingly, we found that the temporal associations of each factor with specific phases of segmentation gene expression are very similar to the temporal associations observed in *Drosophila.* In *Drosophila*, we found that the onset of *prd* expression correlated with the retraction of *cad* expression (Figure 3), and that the early, pair-rule phase of *prd* expression involved regulation by Dichaete (Figure 5); in *Tribolium, Tc-prd* turns on at the anterior limit of the *Tc-cad* domain (Figure 8A-C; Figure S7), with the pair-rule phase of *Tc-prd* expression falling within the *Tc-Dichaete* domain, and stripe splitting occurring anterior to it (Figure 8D-F; Figure S8). In *Drosophila*, we found that the phase 2 (zebra element mediated) expression of *runt* seemed to be activated by Dichaete (Figure 6); in *Tribolium*, the *Tc-runt* pair-rule stripes turn on at the very posterior of the *Tc-Dichaete* domain (Figure 8G-I; Figure S9). Finally, in *Drosophila*, Opa is required for the frequency doubling of pair-rule gene expression and the activation of segment-polarity gene expression (Clark & Akam 2016a; Benedyk et al. 1994); in *Tribolium*, frequency doubling of all the pair-rule genes examined occurs within the *Tc-opa* domain, and the stripes of the segment-polarity genes *Tc-en* and *Tc-wg* emerge within it, as well (Figure 8j-R; Figure S10).

The overall temporal progression of the segmentation process thus seems to be remarkably similar in both species: primary pair-rule genes are expressed dynamically in the context of Cad and Dichaete expression, secondary pair-rule genes turn on as Cad turns off, and frequency doubling and segment-polarity activation occur in the context of Opa expression. Although we have not yet tested these relationships functionally in *Tribolium*, close spatiotemporal associations between segmentation gene expression and our candidate timing factors are evident throughout development, strongly suggesting that patterns of Tc-Cad, Tc-Dichaete, and Tc-Opa activity coordinate the segmentation process, as they do in *Drosophila.* Furthermore, the similarity of the correlations with segmentation gene expression to those in *Drosophila* make it likely that orthologs of Caudal, Dichaete, and Odd-paired temporally regulate the segmentation process in similar (although not necessarily identical) ways in the two different insects.

However, while the regulatory functions of these transcription factors might be broadly conserved between *Drosophila* and *Tribolium*, the spatiotemporal deployment of those functions is obviously divergent. In *Drosophila*, the factors are expressed ubiquitously throughout the trunk and each turns on or off all at once (Figure 2), consistent with controlling the temporal progression of a simultaneous segmentation process. In *Tribolium*, their expression domains are staggered in space, with anterior regions always subjected to a “later” regulatory signature than more posterior tissue (Figure 7). Furthermore, these expression domains retract over the course of germband extension, effectively generating smooth wavefronts that sweep posteriorly across the tissue, consistent with controlling the temporal progression of a sequential segmentation process built around a segmentation clock.

## Discussion

### Caudal, Dichaete, and Odd-paired play temporal regulatory roles in *Drosophila* segmentation

Our dissection of the temporal regulatory roles of Cad and Dichaete builds on other recent work on the *Drosophila* pair-rule gene network (Clark & Akam 2016a; Clark 2017), and represents an important step towards a full understanding of this paradigmatic developmental patterning system. Our findings highlight the overall complexity of pair-rule patterning, which consists of several distinct phases of gene expression, each requiring specific regulatory logic. However, our results also demonstrate how the whole process is orchestrated using just a small number of key regulators expressed sequentially over time. (Although note that there may well be other important timing factors involved in regulating the segmentation process, whose roles are yet to be discovered.) In particular, by rewiring the regulatory connections between other genes, factors like Dichaete and Opa allow a small set of pair-rule factors to carry out multiple different roles. This kind of control logic makes for a flexible and fairly modular regulatory network, and may therefore turn out to be a hallmark of other complex patterning systems.

Further work will be required in order to dissect how this combinatorial regulation is implemented at the molecular level. Dichaete is known to act both as a repressive cofactor (Zhao & Skeath 2002; Zhao et al. 2007) and as a transcriptional activator (Aleksic et al. 2013), therefore a number of different mechanisms are plausible. The Odd-paired protein is also likely to possess both these kinds of regulatory activities (Ali et al. 2012). It should prove particularly interesting to analyse the enhancer regions of genes such as *runt* and prd, which are regulated by both Dichaete and Opa, to find out how exactly their complex expression profiles are generated.

### The segmentation roles of Caudal, Dichaete, and Odd-paired are likely to be broadly conserved between long-germ and short-germ insects

In the second half of this paper, we revealed that Cad, Dichaete, and Opa are likely to play a similar set of roles in *Tribolium* and *Drosophila* segmentation. Given the phylogenetic distance between beetles and flies (separated by at least 300 million years, Wolfe et al. 2016), we suggest that the similarities seen between *Drosophila* and *Tribolium* segmentation are likely to hold true for other insects, and perhaps for many non-insect arthropods as well. These predictions can be tested by comparative studies in various arthropod model organisms. In particular, we anticipate that the expression of cad, *Dichaete*, and *opa* orthologs should retain broadly the same relationship to segmentation gene expression across a wide diversity of developmental modes, and that manipulating the expression of these factors should alter segmentation gene expression accordingly. Note that such manipulations would have to be designed carefully in order to be informative, since *cad* knockdowns in short-germ arthropods cause severe axis truncations (Copf et al. 2004), while Dichaete might exhibit redundancy with other Sox factors, such as SoxN (Overton et al. 2002).

Assuming the temporal regulatory roles of these factors are conserved, they would represent a common spatiotemporal framework for insect segmentation that unifies long-germ and short-germ modes of development. Our findings therefore strengthen recent proposals of regulatory homology between the *Drosophila* early pair-rule gene network and the “segmentation clock” of short germ insects (Clark 2017). This framework could also make sense of other differences between long-germ and short-germ development: for example, if our timing factors turn out to temporally regulate Toll gene expression, the various locations of their expression domains would likely explain the different spatiotemporal distributions of intercalation-based convergent extension observed in different insect species (Paré et al. 2014; Benton et al. 2016)

### The evolution of arthropod segmentation

Finally, we think that the regulatory roles we have uncovered for Cad, Dichaete, and Odd-paired may be highly significant regarding the evolutionary origins and subsequent modification of arthropod segmentation. The patterns of *Tc-cad, Tc-Dichaete*, and *Tc-opa* expression that we have characterised are striking, because essentially similar spatiotemporal expression patterns are observed for these genes in other bilaterian clades, including vertebrates. Cdx genes are expressed in the posterior of vertebrate embryos, where they play crucial roles in axial extension and Hox gene regulation (van Rooijen et al. 2012; Neijts et al. 2016). Sox2 (a Dichaete ortholog) has conserved expression in the nervous system, but is also expressed in a posterior domain, where it is a key determinant of neuromesodermal progenitor (posterior stem cell) fate (Wood & Episkopou 1999; Wymeersch et al. 2016). Finally, Zic2 and Zic3 (Opa orthologs) are expressed in presomitic mesoderm and nascent somites, and have been functionally implicated in somitogenesis (Inoue et al. 2007). All three factors thus have important functions in posterior growth, roles which may well be conserved across Bilateria (Copf et al. 2004).

In *Tribolium*, we think it likely that all three factors are integrated into an ancient gene regulatory network downstream of Wnt signalling, which generates their sequential expression and helps regulate posterior proliferation and/or differentiation (McGregor et al. 2009; Oberhofer et al. 2014; Williams & Nagy 2016). For example, it is suggestive that we observe mutually exclusive patterns of *Tc-wg* and *Tc-Dichaete* in the growth zone: Wnt signalling and Sox gene expression are known to interact in many developmental contexts (Kormish et al. 2010), and these interactions may form parts of temporal cascades (Agathocleous et al. 2009).

We therefore suggest the following outline as a plausible scenario for the evolution of arthropod segmentation. 1) In non-segmented bilaterian ancestors of the arthropods, Cad, Dichaete and Opa were expressed broadly similarly to how they are expressed in *Tribolium* today, owing to conserved roles in posterior elongation. 2) At some point, segmentation genes started to be expressed under the regulatory control of these factors, which represented a preexisting source of spatiotemporal information. Pair-rule genes started oscillating in the posterior, perhaps under the control of Cad (El-Sherif et al. 2014; Schönauer et al. 2016), while the retracting expression boundaries of the timing factors provided wavefronts that effectively translated these oscillations into a periodic patterning of the AP axis, analogous to the roles of the opposing retinoic acid and FGF gradients in vertebrate somitogenesis (Oates et al. 2012). 3) Much later, in certain lineages of holometabolous insects, the transition to long-germ segmentation occurred. This would have involved two main modifications of the short-germ segmentation process: A) changes to the expression of the timing factors, away from the situation seen in *Tribolium*, and towards the situation seen in *Drosophila* (thus causing a major heterochronic shift in the deployment of the segmentation machinery, plus a coordinated shift in all the other processes downstream of the timing factors), and B) recruitment of gap genes to pattern pair-rule stripes, via the ad hoc evolution of stripe-specific elements. This transition process would likely have occurred gradually along the body axis, from anterior to posterior (Peel & Akam 2003; Rosenberg et al. 2014).

In other words, this model involves first creating new regulatory interactions between existing domains of expression and the downstream genes that are to be patterned, and later changing these upstream expression domains, without altering the existing downstream regulatory network. The recruitment/reuse of existing developmental features at each stage reduces the number of regulatory changes that would have to evolve de novo, facilitating the evolutionary process. Note that under this hypothesis arthropod segmentation would not be homologous to segmentation in other phyla, but would probably have been fashioned from common parts (Chipman 2010; Graham et al. 2014).

## METHODS

### Drosophila melanogaster

Embryo collections were carried out at 25°C. The *Drosophila* mutants used were cad^2^ (gift of Helen Skaer), D^r72^ (gift of Steve Russell), eve^3^ (gift of Bénédicte Sanson), and h^22^ (Bloomington stock no. 5529). Wild-type flies were Oregon-R. In order to distinguish homozygous mutant embryos, mutant alleles were balanced over *CyO hb::lacZ* (Bloomington stock no. 6650) or *TM6c Sb Tb twi::lacZ* (Bloomington stock no. 7251). DIG-labelled and FITC-labelled riboprobes were generated using full-length cDNAs from the *Drosophila* gene collection (Stapleton et al. 2002), obtained from the *Drosophila* Genomics Resources Centre. The clones used were LD29596 (cad), LD16125 (opa), RE40955 (hairy), MIP30861 (eve), IP01266 (runt), GH22686 *(ftz)*, RE48009 (odd), GH04704 (prd), LD30441 (slp), and F107617 (en). The cDNA for *Dichaete* was a gift from Steve Russell, and the cDNA for *lacZ* was a gift from Nan Hu.

Double fluorescent in situ hybridisation was carried out as described previously (Clark & Akam 2016a). Images were acquired using a Leica SP5 confocal microscope. Contrast and brightness adjustments of images were carried out using Fiji (Schindelin et al. 2012; Schneider et al. 2012). Some of the wild-type images were taken from a previously published dataset (Clark & Akam 2016b).

### Tribolium castaneum

*Tribolium castaneum* eggs were collected on organic plain white flour (Doves Farm Foods Ltd, Hungerford, U.K.) at 30°C over a period of 48 hours. Alkaline phosphatase in situ hybridization on whole mount embryos were carried out as previously described (Schinko et al. 2009). RNA probes were DIG-labelled (all genes) and in most cases also FITC-labelled *(Tc-Dichaete, Tc-opa, Tc-prd, Tc-wg* & *Tc-en)* and prepared according to Kosman et al. (2004), using gene fragments amplified from *Tribolium castaneum* genomic DNA (for *Tc-cad, Tc-eve, Tc-odd* & *Tc-run)* or cDNA (for *Tc-Dichaete, Tc-opa, Tc-prd, Tc-wg* & *Tc-en)* and cloned into the pGEM-T Easy Vector (Promega, Madison, WI). Clones, and their details, are available on request.

The generation of DIG-labelled probes against *Tc-cad* has been previously described in Peel & Averof (2010), and the generation of DIG-labelled probes against *Tc-eve* and *Tc-odd* has been previously described in (Sarrazin et al. 2012). The remaining gene fragments were amplified using the following primers: *Tc-run:* 5’-CAACAAGAGCCTGCCCATC-3’ & 5’-TACGGCCTCCACACACTTT-3’ (amplifies 3,158bp fragment). *Tc-Dichaete* (TC013163): 5’-TAACAACCGACACCCAACAG-3’ & 5’-TTGACGACCACAGCGATAATAA-3’ (921bp fragment). *Tc-opa* (TC010234): 5’-CCCAAGAATGGCCTACTGC-3’ & 5’-TTGAAGGGCCTCCCGTT-3’ (710bp 5’ fragment), 5’-GCGAGAAGCCGTTCAAAT-3’ & 5’-TCTCTTTATACAATTGTGGTCCTAC-3’ (705bp 3’ fragment); two probes made separately and combined. *Tc-prd:* 5’-GAATACGGCCCTGTGTTATCT-3’ & 5’-ACCCATAGTACGGCTGATGT-3’ (1179bp fragment). *Tc-wg:* 5’-CAACGCCAGAAGCAAGAAC-3’ & 5’-ACGACTTCCTGGGTACGATA-3’ (1095bp fragment). *Tc-en:* 5’-TGCAAGTGGCTGAGTGT-3’ & 5’-GCAACTACGAGATTTGCCTTC-3’ (1001bp fragment).

In the double in situ hybridizations where *Tc-cad* mRNA is detected in red, the primary antibodies were switched such that anti-DIG-AP was used second (after anti-FITC-AP) to detect *Tc-cad DIG* probe and signal developed using INT/BCIP (see Schinko et al. (2009) for more details). Embryos were imaged on a Leica M165FC Fluorescence Stereo Microscope with a Q Imaging Retiga EXI colour cooled fluorescence camera and Q Capture Pro 7 software.

## ACKNOWLEDGEMENTS

EC would like to thank Michael Akam and Tim Weil for advice, lab resources and encouragement. ADP would like to thank Ian Hope and Stephanie Wright for the use of their lab equipment. EC & ADP would like to thank Michael Akam and Toby Andrews for insightful comments on the manuscript.

---

**Fig. S1:**
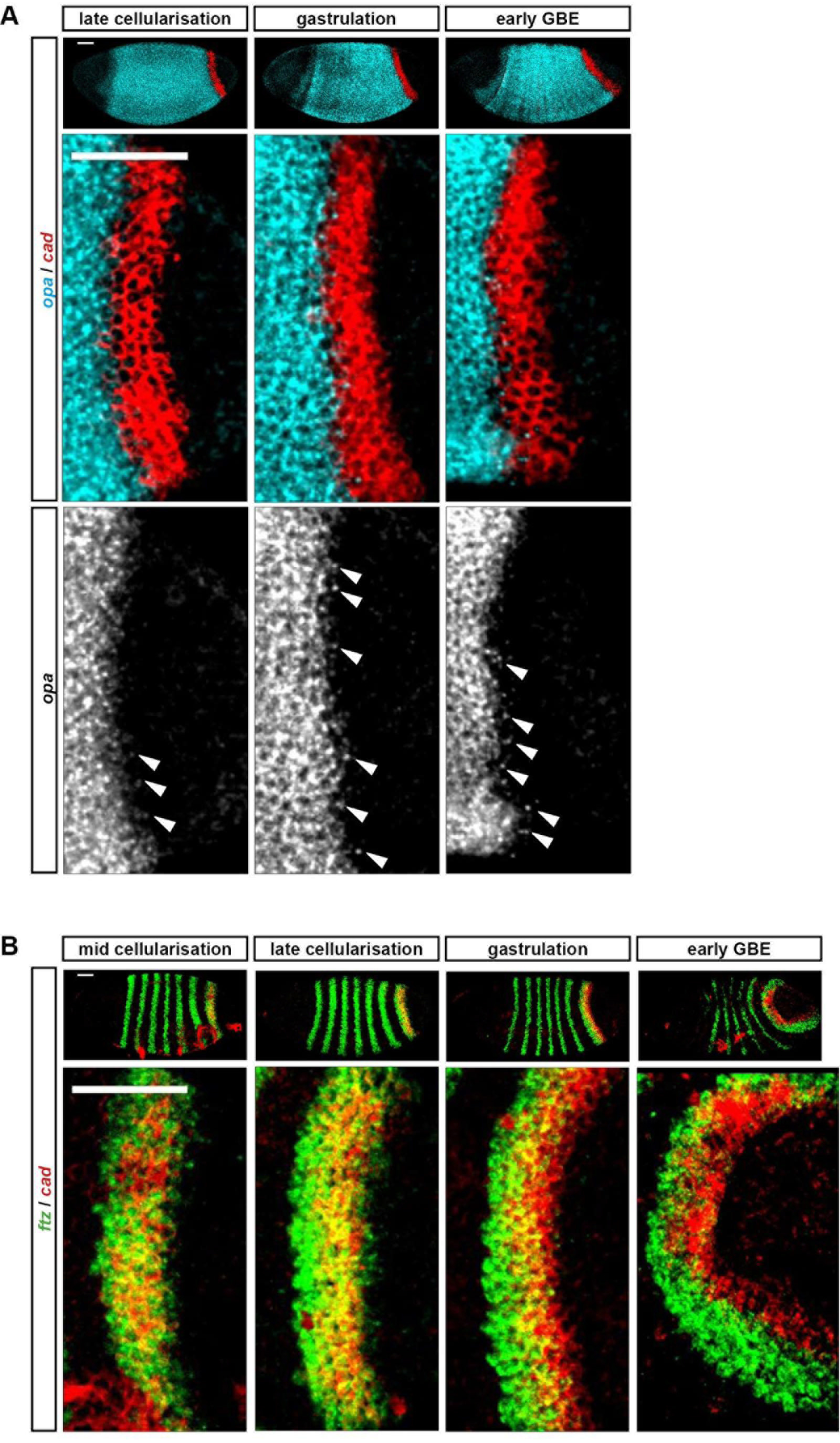
Shifting boundaries in the posterior of the embryo. **A:** Nascent transcription (nuclear dots, marked by arrowheas) of *opa* expression are observed within the *cad* domain, indicative that the *opa* expression domain is expanding posteriorly. Scale bars = 50 μm. **B:** Marked posterior shifting of the *cad* posterior domain relative to stripe 7 of*ftz.* Cropped and rotated enlargements of the stripe 7 region are show below the whole embryo views. (The bright red regions of staining outside the *cad* posterior domain are artefacts caused by bits of debris stuck to the embryos.) Scale bars = 50 μm. Note that without live reporters, it is hard to determine unambiguously which of the cad, *Dichaete, opa*, and *odd* expression domains are moving relative to cells, and which are static (see Figure 2D,E). Nevertheless, we are reasonably sure that the cad, *Dichaete*, and *opa* domains are dynamic, for three reasons. First, quantitative measurements in populations of fixed embryos indicate that, in contrast to other primary pair-rule stripes, neither *odd* stripe 7 nor *ftz* stripe 7 **(B)** shift significantly within the embryo (Surkova et al. 2008). Second, *odd* stripe 7 is repressed anteriorly by Opa expression, since it emerges within Runt stripe 7, and Runt+Opa combined repress *odd* (Clark & Akam 2016a). Third, at the same time as the domains are shifting relative to one another, the *opa* domain appears to be expanding posteriorly **(A).** These anterior-to-posterior shifts of cad, *Dichaete*, and *opa* expression are a notable counterpoint to the posterior-to-anterior shifts observed for gap and pair-rule gene expression (Jaeger et al. 2004; Surkova et al. 2008), and it will be interesting to find out how they are generated.

**Fig. S2:**
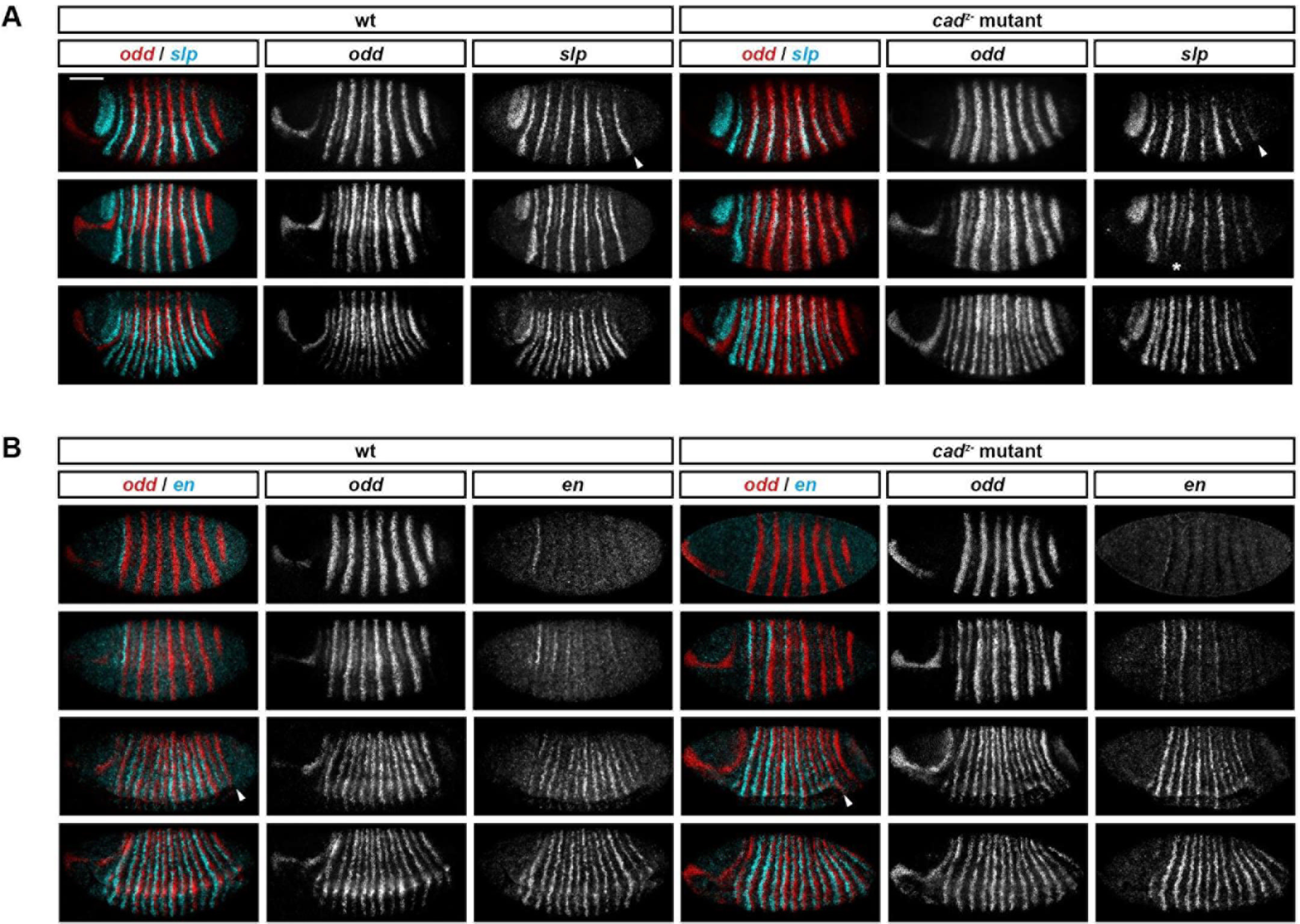
Normal temporal patterning of *slp, odd*, and *en* in *cad* zygotic mutants. A: *slp* expression is emerges at the normal time in *cad* zygotic mutants, although there are some changes to the intensity of the stripes, particular in anterior ventral regions (asterisk). DV patterning of the *slp* stripes is normal. Arrowheads mark the emergence of *slp* stripe 7, which is considerably weakened in *cad* zygotic mutants, likely owing to the posterior expansion of the fate map (by about 10% AP length, based on measurements of posterior stripes). In the mutant, this stripe probably experiences a different background of activators and repressors than the more anterior stripes, which are located within the main trunk. Scale bar = 100 μm. **B:** *en* expression emerges at the correct time in *cad* zygotic mutants, although the order of stripe emergence and the DV patterning of the stripes is altered. Arrowheads mark the posterior extent of the ventral furrow, which aligns with the posterior extent of segmentation gene expression in wild-type embryos, but not *cad* zygotic mutants.

**Fig. S3:**
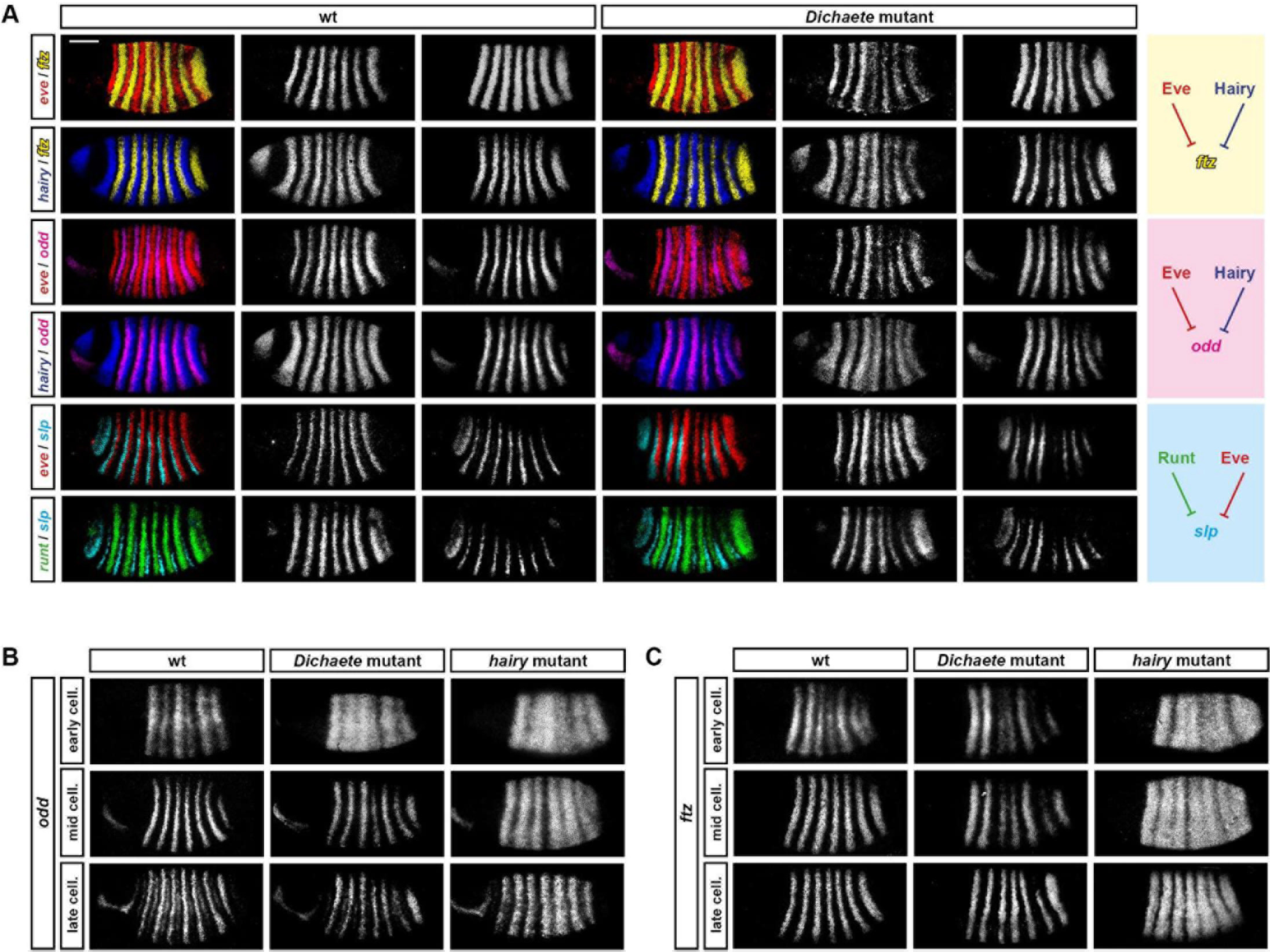
*ftz, odd*, and *slp* are cross-regulated normally in *Dichaete* mutant embryos. **A:** Expression of*ftz, odd*, and slp, relative to their pair-rule repressors, at mid-cellularisation (phase 2). Single channel images are shown to the right of each false-coloured dual-channel image *(ftz, odd*, and *slp* are always on the right). Right: arrow diagrams describing ftz, odd, and *slp* regulation during phase 2. Hammerhead arrows represent repression. Activators (likely to be ubiquitously-expressed factors) are not included. Scale bar = 100 μm. **B, C:** Expression of *odd* **(B)** or *ftz* **(C)** in wild-type, *Dichaete* mutant, and *hairy* mutant embryos, over the course of cellularisation. Note the ectopic *odd* expression seen at early cellularisation in *Dichaete* mutants.

**Fig. S4:**
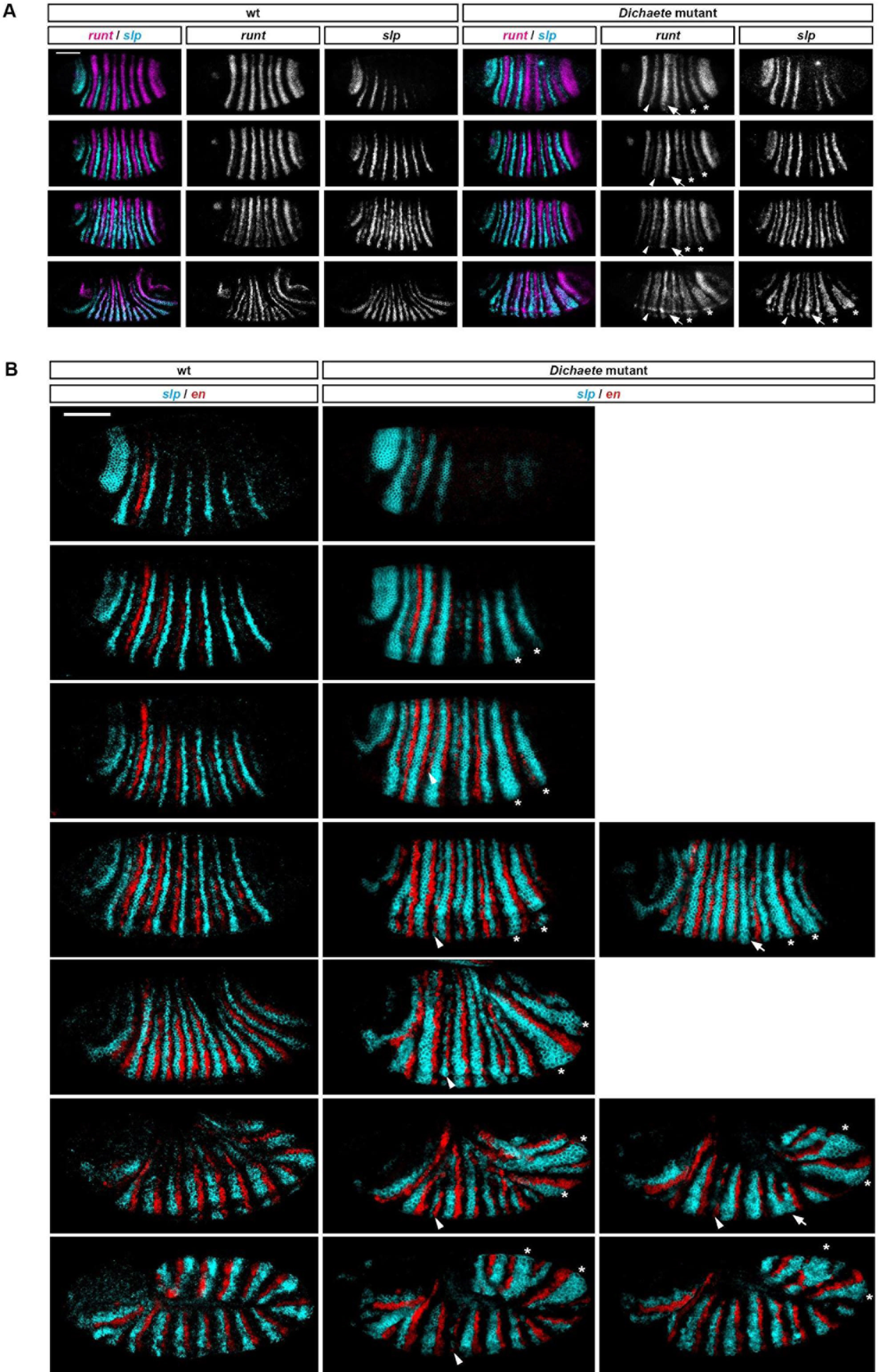
Late pair-rule gene expression in *Dichaete* mutants reveals the aetiology of the *Dichaete* cuticle phenotype. **A:** Irregular *runt* expression in *Dichaete* mutants at cellularisation (top) results in an irregular pattern of *slp* expression at the end of gastrulation (bottom). Embryo age increases from top to bottom. Asterisks indicate weakened *runt* stripes 5 and 6 in the mutant embryos. These stripes are responsible for patterning the anterior boundaries of *slp* stripes 6 and 7, which tend to be expanded anteriorly in the mutants. (Note that the third embryo down is unusually mildly affected, compare equivalently-stage *slp* expression in **B.)** These anterior expansions often result in fusions with the fifth and sixth secondary *slp* stripes (asterisks in bottom row, right). Late *runt* is patterned in the same way as slp, meaning that the fifth and sixth *runt* secondary stripes form contiguously with the posteriors of the primary stripes (asterisks in bottom row, left). Arrowheads in the mutant point to weakened *runt* stripe 2, which leads to an expanded *slp* stripe 3, and eventually a narrow and incomplete second *slp* secondary stripe (arrowhead in bottom row, right). Finally, arrows in the mutant point to the expanded *runt* stripe 3, which represses *slp* stripe 4 at early stages, and can lead to patterning irregularities later on (arrow, bottom row, right). Scale bar = 100 μm. **B:** Irregular boundary specification in *Dichaete* mutant embryos. Parasegment boundaries are specified in wild-type embryos (left) by abutting stripes of *slp* and *en* expression, separated by clear buffer zones (correspond to segmental stripes of *odd* expression), which allow the boundaries to maintain polarity. The irregular patterns of *slp* expression, resulting ultimately from irregular *runt* expression (see **A)** lead to various boundary specification defects: fusions with more anterior boundaries (asterisks); narrow/incomplete boundaries (arrowheads); and fusions with more posterior boundaries (arrow). Scale bar = 100 μm.

**Fig. S5.**
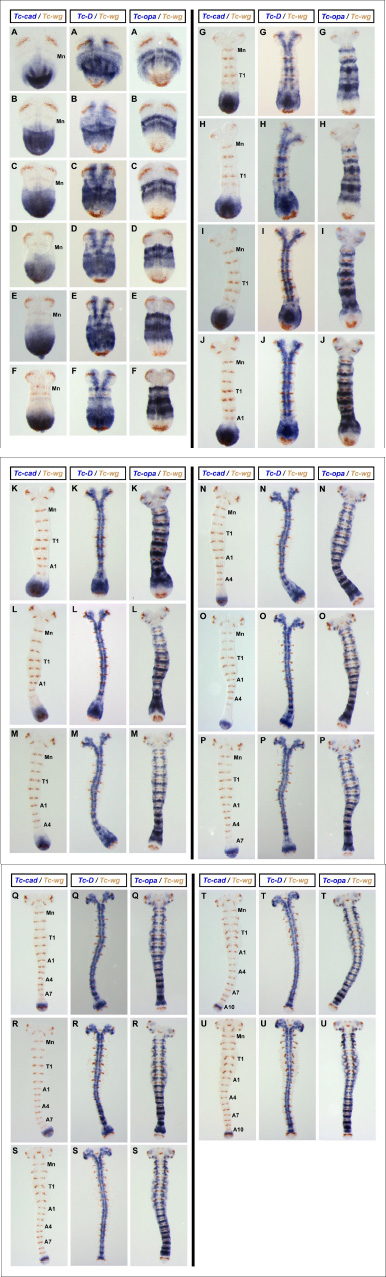
Expression of *Tc-cad, Tc-Dichaete* & *Tc-opa* relative to a common segment marker, *Tc-wg*, in *Tribolium castaneum* germband stage embryos. **(A-U)**. Sets of three *Tribolium castaneum* germband stage embryos that have been stage matched using *Tc-wg* expression patterns *(Tc-wg* expression brown in all panels). Stage matched germband embryos increase in age from **A** to **U.** In each set of embryos, the left-hand embryo is also stained for *Tc-cad* expression, the middle embryo is stained for *Tc-Dichaete* expression and the right-hand embryo is stained for *Tc-opa* expression (all blue stains). In the double in situ hybridizations for *Tc-cad* & *Tc-wg* (left-hand embryos) the mandibular (Mn), prothoracic (T1), 1^st^ abdominal (A1), 4^th^ abdominal (A4), 7th abdominal (A7) and/or 10^th^ abdominal (A10) stripes of *Tc-wg* expression have been labeled. Note how the relative position of the expression domains of these three genes is remarkably conserved across progressive germband elongation stages. Consult Fig. 7 for a clear comparison across different developmental stages, rather than between *Tc-cad, Tc-Dichaete* & *Tc-opa* expression patterns.

**Fig. S6.**
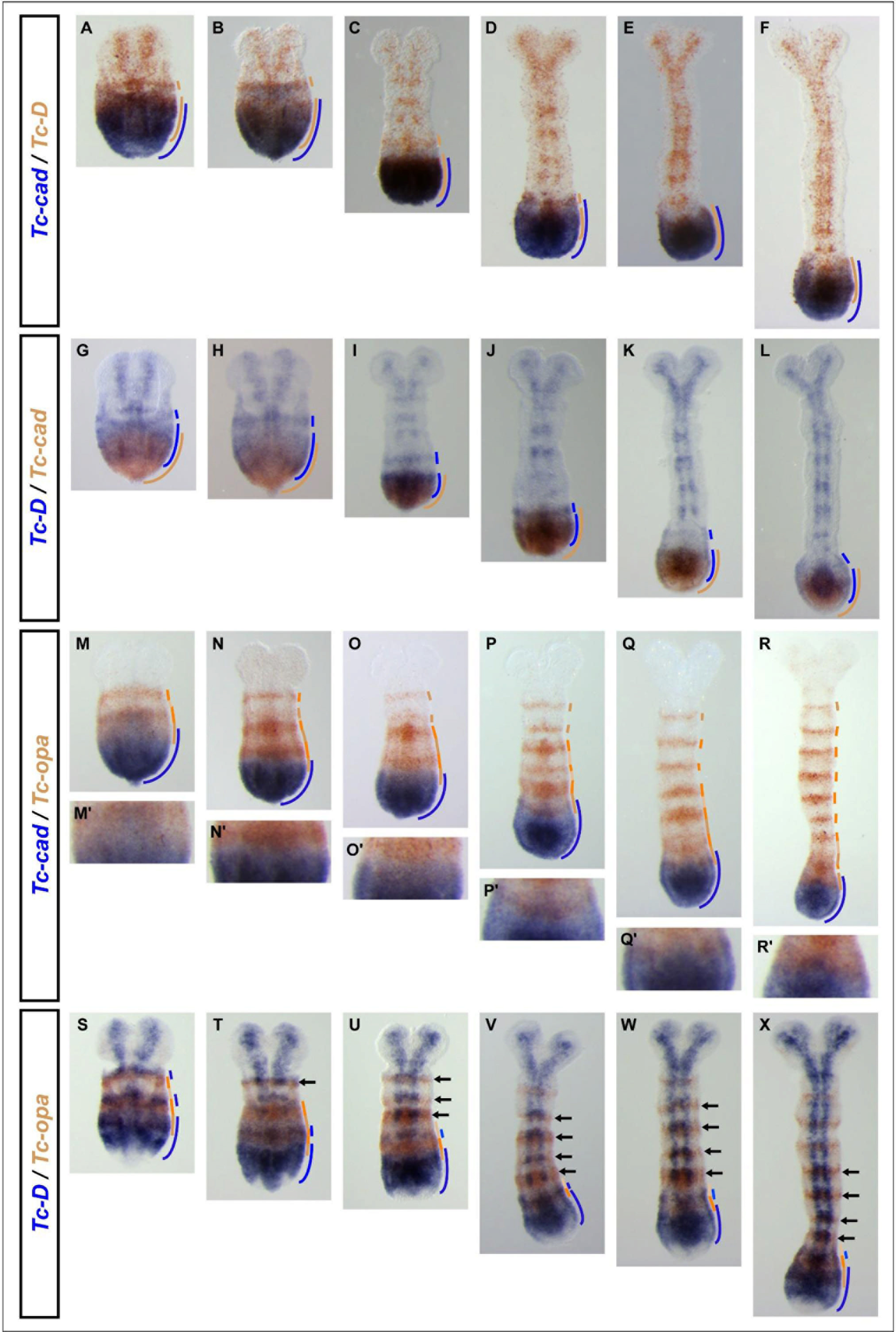
Expression patterns of *Tc-cad, Tc-Dichaete* & *Tc-opa* in relation to each other in *Tribolium castaneum* germband stage embryos. **(A-F).** Double in situ hybridization for *Tc-cad* (blue) and *Tc-Dichaete* (brown) in embryos of increasing age from left **(A)** to right **(F). (G-L).** As for panels **A-F,** but this time *Tc-Dichaete* DIG and *Tc-cad* FITC RNA probes were used instead of *Tc-cad* DIG and *Tc-Dichaete* FITC RNA probes such that the colours are reversed. Note the stripe of *Tc-Dichaete* expression that is observed anterior to the *Tc-cad* domain in some, but not all, embryos (see text for further details). **(M-R).** Double in situ hybridization for *Tc-cad* (blue) and *Tc-opa* (brown) in embryos of increasing age from left **(M)** to right **(N).** Panels **M’-R’** show higher magnification images of the regions in **M-R** where *Tc-cad* and *Tc-opa* expression overlaps. These data suggest that as posterior germband cells move anteriorly relative to the posterior tip of the elongating embryo due to convergent extension cell movements, they experience a drop as *Tc-cad* expression levels as *Tc-opa* expression levels increase. **(S-X).** Double in situ hybridization for *Tc-Dichaete* (blue) and *Tc-opa* (brown) in embryos of increasing age from left **(S)** to right **(X).** Black arrows point to late *Tc-opa* segmental stripes that overlap strong, segmentally-reiterated *Tc-Dichaete* expression domains that are limited to the medially positioned neuroectoderm. Colour-coded lines on the right-hand side of the embryos indicate our interpretations of the expression patterns in **A-X.**

**Fig. S7.**
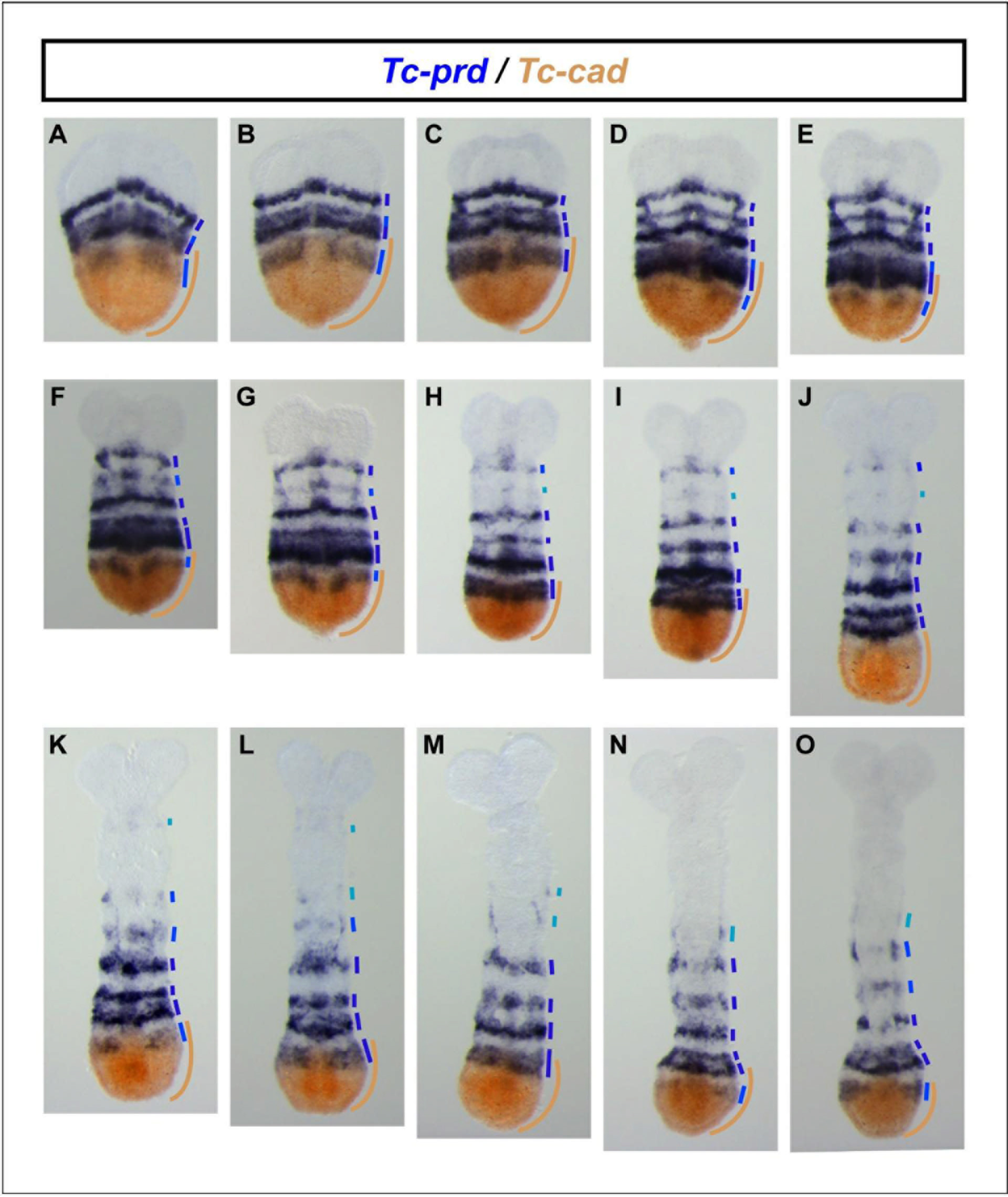
Expression of *Tc-prd* relative to *Tc-cad* in *Tribolium castaneum* germband stage embryos. **(A-O).** Double in situ hybridization for *Tc-prd* (blue) and *Tc-cad* (brown) in embryos of increasing age from youngest **(A)** to oldest **(O).** Colour-coded lines on the right-hand side of the embryos indicate our interpretations of the expression patterns in **A-O.** Note how the primary pair-rule stripes of *Tc-prd* first appear and form within the anterior half of the posterior *Tc-cad* domain (see where blue lines overlap brown lines). In contrast, segmental stripes of *Tc-prd* expression resolve by splitting just anterior to the *Tc-cad* domain (see where blue lines lie anterior to the brown line).

**Fig. S8.**
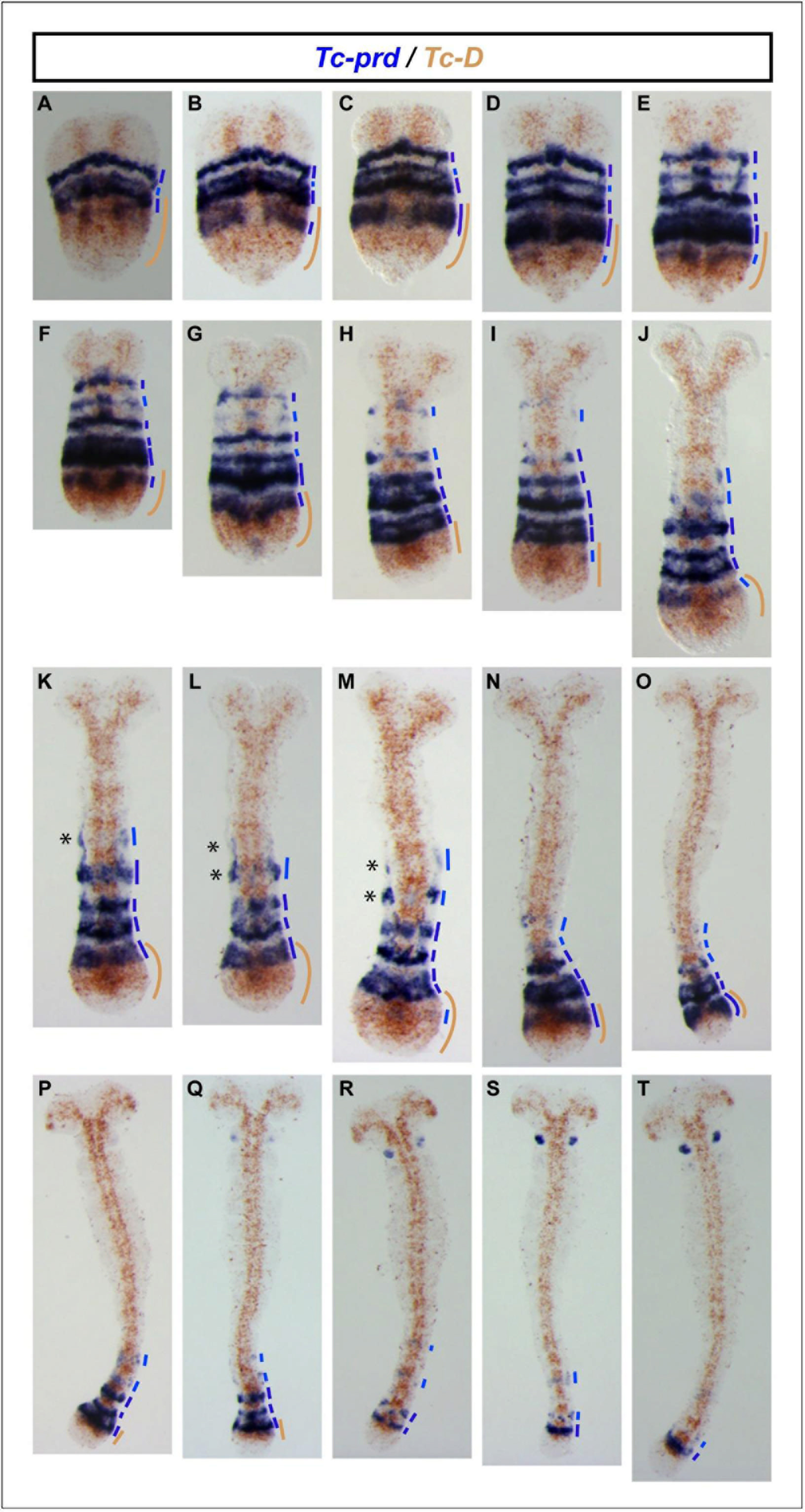
Expression of *Tc-prd* relative to *Tc-Dichaete* in *Tribolium castaneum* germband stage embryos. **(A-T).** Double in situ hybridization for *Tc-prd* (blue) and *Tc-Dichaete* (brown) in embryos of increasing age from youngest **(A)** to oldest **(T).** Colour-coded lines on the right-hand side of the embryos indicate our interpretations of the expression patterns in **A-T.** Note how the primary pair-rule stripes of *Tc-prd* first appear and form within the posterior-most *Tc-Dichaete* domain (see where blue lines overlap brown lines). In contrast, segmental stripes of *Tc-prd* expression resolve by splitting anterior to this domain (see where blue lines lie anterior to the brown line). While dissecting and cleaning the embryos we noted that *Tc-prd* expression remains on stronger and longer in the overlying amnion compared to the underlying ectoderm; this is particularly apparent in panels **K** to **M** where the *Tc-prd* stained amnion has been ripped away while cleaning the embryo of yolk to reveal ectoderm free from *Tc-prd* expression (asterisks). Amnion-related expression can be seen down the lateral margins of many of the embryos where some amniotic cells survived dissection and cleaning.

**Fig. S9.**
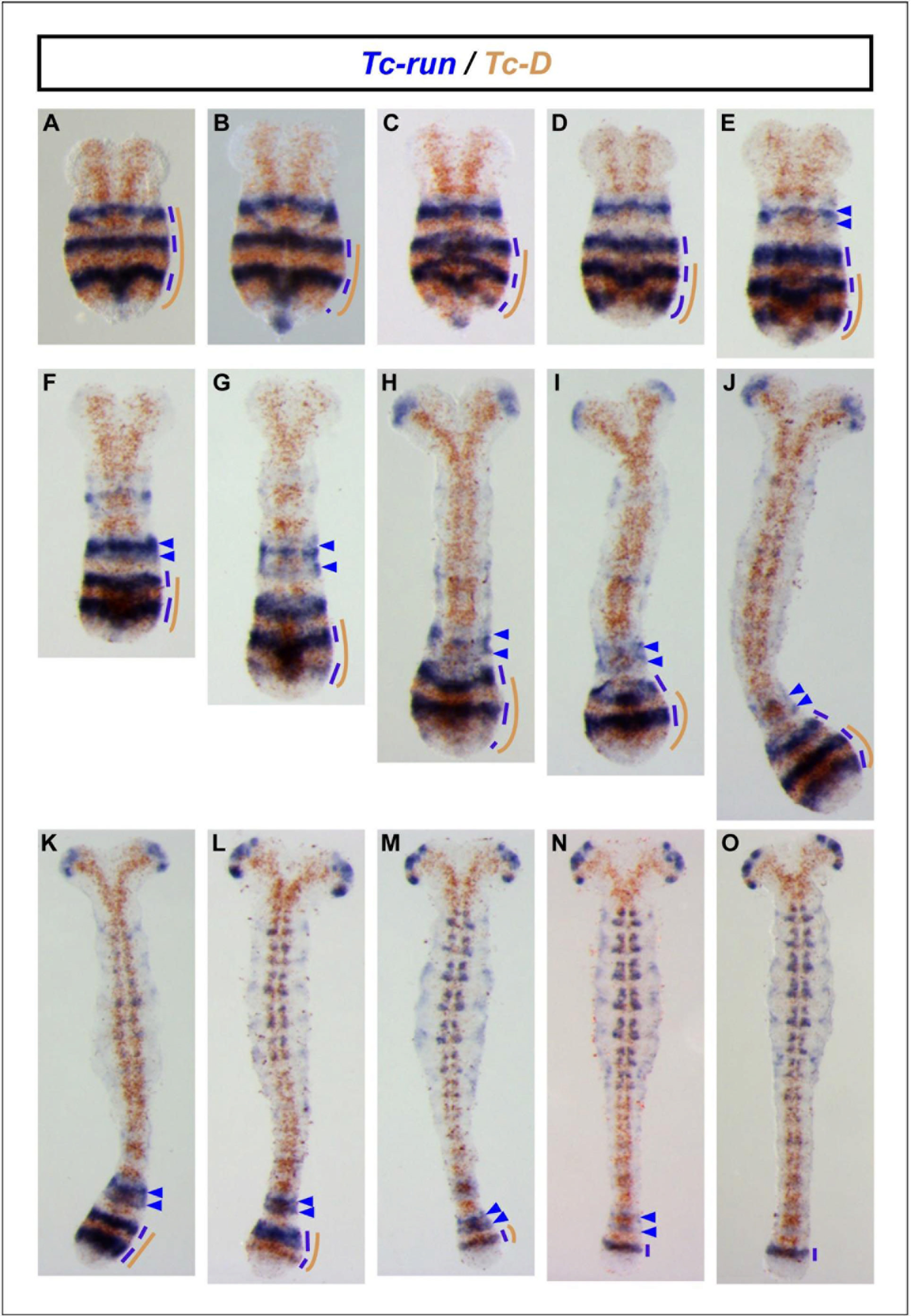
Expression of *Tc-run* relative to *Tc-Dichaete* in *Tribolium castaneum* germband stage embryos. **(A-O).** Double in situ hybridization for *Tc-run* (blue) and *Tc-Dichaete* (brown) in embryos of increasing age from youngest **(A)** to oldest **(O).** Colour-coded lines on the right-hand side of the embryos indicate our interpretations of the expression patterns in **A-O.** In **A-O** blue arrowheads mark the primary pair-rule stripes that have most recently resolved – or are in the process of resolving - to a segmental periodicity. In some younger embryos **(A-H),** more than two stripes are apparent due to differences in the timing and/or positioning of this process between the amnion and ectoderm cell layers. Note how *Tc-run* stripe splitting occurs anterior to the posterior-most *Tc-Dichaete* domain (as judged by brown line by side of embryo). Older embryos show additional domains of *Tc-run* expression in the head lobes **(H-O)** and neuroectoderm **(J-O)** that were used to help stage the embryos.

**Fig. S10.**
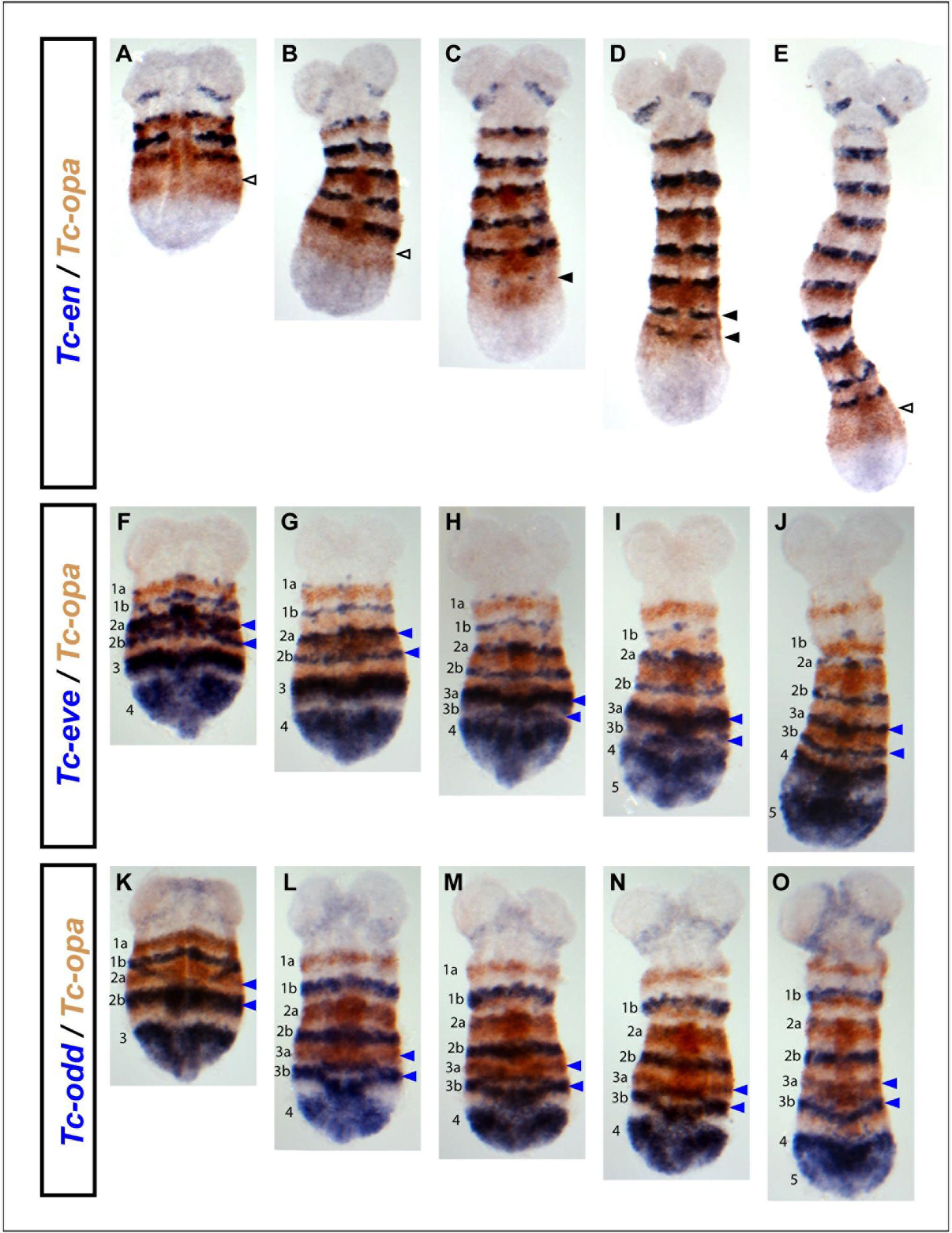
Expression of *Tc-opa* relative to *Tc-en, Tc-eve* and *Tc-odd* in *Tribolium castaneum* germband stage embryos. **(A-E).** Double in situ hybridization for *Tc-en* (blue) and *Tc-opa* (brown) in embryos of increasing age from left **(A)** to right **(E).** Note how the *Tc-en* stripes (solid black arrowheads in **C, D)** form within the *Tc-opa* domain, but in a stripe-shaped region that is already clearing of *Tc-opa* expression (empty black arrowheads in **A, B, E). (F-J).** As for **A-E,** but this time double in situ hybridization for *Tc-eve* (blue) and *Tc-opa* (brown). **(K-O).** As for **A-E,** but this time double in situ hybridization for *Tc-odd* (blue) and *Tc-opa* (brown). In **F-O,** note how the segmental stripes of *Tc-odd* and *Tc-eve* (labeled a & b) resolve within the *Tc-opa* domain. In **F-O,** blue arrowheads mark *Tc-odd* & *Tc-eve* segmental stripes that have most recently resolved, or are in the process of resolving.

